# Oligodendrocyte precursor cells stop sensory axons regenerating into the spinal cord

**DOI:** 10.1101/2022.09.28.509944

**Authors:** Hyukmin Kim, Andy Skuba, Jingsheng Xia, Sung Baek Han, Jinbin Zhai, Huijuan Hu, Shin H. Kang, Young-Jin Son

## Abstract

Primary sensory axons stop regenerating as they re-enter the spinal cord, resulting in incurable sensory loss. What arrests them has remained unclear. We previously showed that axons stop by forming synaptic contacts with unknown non-neuronal cells. Here, we identified these cells in adult mice as oligodendrocyte precursor cells (OPCs). We also found that only a few axons stop regenerating by forming dystrophic endings, exclusively at the CNS:PNS borderline where OPCs are absent. Most axons stop in contacts with a dense network of OPC processes. Live imaging, immuno-EM and OPC-DRG co-culture additionally suggest that axons are rapidly immobilized by forming synapses with OPCs. Genetic OPC ablation enables many axons to continue regenerating deep into the spinal cord. We propose that sensory axons stop regenerating by encountering OPCs that induce presynaptic differentiation. Our findings identify OPCs as a major regenerative barrier that prevents intraspinal restoration of sensory circuits following spinal root injury.

## INTRODUCTION

The dorsal root (DR) carries primary sensory axons projecting centrally from DR ganglia (DRG). DR injuries, which commonly result from brachial plexus trauma, cause incurable loss of sensation, motor incoordination and chronic pain ^1^. The devastating consequences are because DR axons stop regenerating at the entrance of the spinal cord, the dorsal root entry zone (DREZ), and therefore fail to restore connections with intraspinal neurons. Both neuron-intrinsic and - extrinsic mechanisms, which limit axon regrowth elsewhere in the CNS ^2-4^, are believed to prevent re-entry of DR axons. Unlike spinal cord injury, DR cut or crush injury damages axons in the PNS without causing an impassable glia scar at the DREZ. Nevertheless, DR axons stop at the scar-free DREZ, even after a nerve conditioning lesion that boosts regenerative ability of DRG neurons ^5,6^.

How the scar-free DREZ or spinal cord stops axons remains unclear, but myelin-associated inhibitors (MAIs) and chondroitin sulfate proteoglycans (CSPGs) are conventionally considered responsible ^7,8^. Surprisingly, however, co-eliminating MAIs (Nogo/Reticulon-4, MAG, OMgp) and CSPGs enables only a few conditioning-lesioned axons to extend a short distance beyond the DREZ ^9^. These findings led us to speculate that unknown mechanism(s), unrelated to MAIs and CSPGs, may primarily account for the vigorous arrest of DR axons at the DREZ.

Our earlier findings also challenged the notion that DR axons stop by forming dystrophic endings or retraction bulbs ^7^, like severed CNS axons degenerating after spinal cord injury ^10,11^. This prevalent notion was based on the large bulbous endings occasionally reported in light- or ultramicroscopic studies. The first *in vivo* imaging of regenerating DR axons, however, revealed that axons rapidly stop at the DREZ and remain completely immobilized, with only a few enlarging their tips to resemble dystrophic endings. Furthermore, ultrastructural surveys targeting immobilized axons disclosed synapse-like or apparent presynaptic endings that formed on unknown non-neuronal cells ^6^. These findings led us to suggest that unknown non-neuronal cells stop regenerating axons by eliciting presynaptic differentiation ^6,12^.

Here, we report that these non-neuronal, postsynaptic cells are oligodendrocyte precursor cells (OPCs), also known as NG2 glia, that form bona fide synapses with unmyelinated axons widely in the adult CNS ^13-15^. Additionally, we suggest that OPCs may play a more decisive role than other extrinsic mechanisms, and that dystrophic endings form infrequently in the absence of OPC contacts. These findings provide new insights into the regeneration failure of sensory axons at and beyond the DREZ, and the pathophysiological significance of OPCs in the traumatized nervous system.

## RESULTS

### Only a few axons stop by forming dystrophic endings exclusively at the borderline

Most DR axons do not stop regenerating at the CNS:PNS glia borderline where astrocytes/oligodendrocytes are tightly juxtaposed to Schwann cells. They cross the borderline and then stop at the DREZ, which is a region of CNS tissue that extends for ∼100μm beyond the borderline (see Figure 3C) ^6,9^. Because we did not observe retraction bulbs or dystrophic endings, we extended our earlier work by expanding ultrastructural surveys to look for the formation of these endings in a broader area of the DREZ.

We crushed cervical roots of adult mice and prepared serial longitudinal slices of dorsal spinal cord containing a root at 2 wpi. These preparations facilitated distinguishing CNS and PNS territory across the borderline under electron microscopy (EM; e.g., Figure 5C). Consistent with our earlier work, we frequently observed synapse-like contacts at the DREZ that appeared to form on non-neuronal cells (Figure 1A). In contrast, thorough searches of wide areas on numerous slices only occasionally revealed large bulbous, disorganized axon endings. These endings were 3 to 5 times larger than axons and densely filled with disorganized multi-organelles and scattered vacuoles (Figure 1B). Notably, dystrophic endings were usually observed at the borderline, but not in deeper CNS territory of the DREZ. Furthermore, unlike synapse-like contacts, these large endings were embedded in collagen fibers and displayed a noteworthy absence of cellular contacts (Figure 1B). Similarly, we noted that dystrophic endings described by previous authors also lacked cellular contacts and were observed in PNS territory beyond the DREZ ^5^.

**Figure 1.**
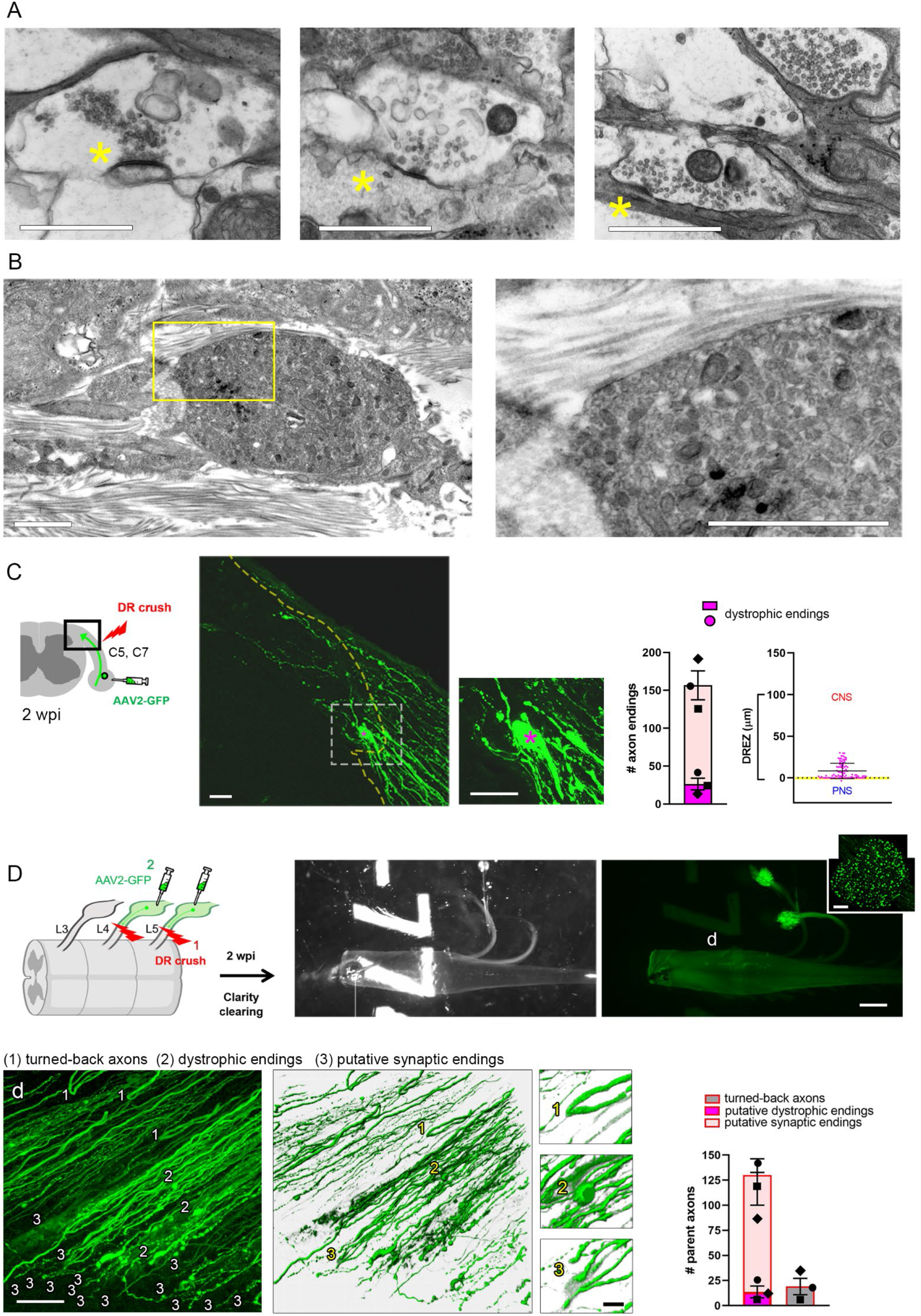
Dystrophic endings form by axons embedded in collagen fibers at the borderline. **(A, B)** Ultrastructural comparison of synaptic-like profiles (A) and a dystrophic ending (B) formed by DR axons at the DREZ at 2 wpi. The presynaptic axons exhibiting clustered vesicles are small and opposed to unknown postsynaptic cells exhibiting thin postsynaptic density (asterisks), whereas the dystrophic ending is large, disorganized, and embedded in acellular collagen fibers. Scale bar: 500nm **(C)** The incidence and location of dystrophic endings analyzed on transverse sections, at 2 weeks after C5-C8 root crush followed by AAV2-GFP injections to C5 and C7 DRGs. A representative section, with enlarged view, showing a large dystrophic ending (asterisk) formed at the borderline (dotted line). Note that many other axons stopped past the borderline do not display dystrophic endings. Graphs show the incidence of dystrophic endings and their location relative to the borderline (n= >100 axons/mouse, 3 mice; putative synaptic endings, 156.7±19.10; dystrophic endings, 26.33±7.688; locations of dystrophic endings (n=91), 8.308±9.233). Error bars represent SD. Scale bar: 20μm **(D)** High resolution 3D imaging and quantitative analysis of axon endings on wholemounts of CLARITY cleared spinal cords at 2 weeks after L4 and L5 root crush. Transmitted light and fluorescent views of a representative cleared spinal cord, roots, DRG (inset), and GFP+ axons arrested at the DREZ in enlarged (d) or volume rendered images. Representative examples illustrate parental axons that turn around (e.g., 1), form a large dystrophic ending (e.g., 2), or form a small putative synaptic ending (e.g., 3). Relative incidence of different axon endings illustrating that a limited number of parental axons turned around or stopped by forming dystrophic endings. n= >100 axons/mouse, 3 mice. Putative synaptic endings, 116.3±16.33; turned-back axons, 13.67±5.925; dystrophic endings, 19.0±8.185. Error bars represent SD. Scale bars: 200μm (clearing), 50μm (DRG), 50μm (3D, volume), 10μm (enlarged 3D)

We next employed light microscopic approaches to examine the incidence and location of dystrophic endings. In the first, we selectively labeled regenerating axons by microinjecting AAV2-GFP into DRGs after crushing cervical roots and assessed them in serial transverse sections at 2 wpi (Figure 1C). Some sections displayed large bulbous, presumptive dystrophic, endings, which represented ∼5% axons. Consistent with ultrastructural surveys, these endings were observed close to the borderline, but not deep in the CNS territory. Our second approach was to examine DR axons in their entirety without sectioning. To do this, we performed Clarity tissue clearing at 2 wpi, and examined fluorescently labeled axons in an entire root prepared in wholemounts (Figure 1D). High resolution 3D imaging of cleared roots revealed a small number of parent axons that turned around or formed dystrophic endings near the borderline. These axons represented ∼15% and ∼10% of all GFP+ axons, respectively. In contrast, most axons extended further and then stopped by forming small endings and frequent swellings on their shafts resembling axonal varicosities. Collectively, together with our earlier data, these findings show that DR axons stop by forming synapse-like contacts or dystrophic endings. They also suggest that dystrophic endings are formed at the borderline by occasional axons entrapped in collagen fibers, unlike synapse-like contacts formed by many other axons making cellular contacts after crossing the borderline.

### Two-photon imaging confirms rapid axon immobilization at the DREZ

In contrast to the prevailing view, these data suggest that only a few axons stop by forming dystrophic endings. The conventional view was primarily based on the observations that DRG axons *in vitro* retract in response to MAIs ^16^, or stall and become dystrophic when exposed to a gradient of CSPGs ^17^. Moreover, these dystrophic endings remain extremely motile at least for several days ^17,18^. In contrast, DR axons that we monitored in living mice with widefield microscopy did not retract, but were rapidly immobilized as they entered the DREZ ^6^. To ensure that we have not overlooked subtle axon motility, we performed live imaging using high-resolution two-photon microscopy.

For optimal two-photon imaging, it is essential that fluorescent labeling mark only a small number of DR axons ^19^. Accordingly, we used the M line of *Thy1* mice (Thy1-M), in which <5% of DRG neurons are YFP+ ^20^. Following DR crush, axons penetrate the crush site in a day or two, and then regenerate along the root at a speed of ∼1 mm/day ^6^. We crushed an L5 root >4 mm from the borderline (Figure 2A), to make regenerating axons arrive at the DREZ as early as 5 days post-crush. We then screened ∼150 mice on the first day of imaging at 4 or 5 dpi, to identify ∼20 mice with 1 (or rarely 2) YFP+ axons superficially positioned within the optimum imaging depth (∼150μm below the dura; Figure 2B). We then continuously monitored the distal endings of these axons for up to 5 hours. As anticipated, we observed axons at the DREZ at 5 dpi, but none at 4 dpi. Notably, no axons moved forward or retracted; all appeared already stopped at the DREZ. Axon tips with either a slender (Figure 2C) or small roundish shape (Figure 2D) exhibited no perceptible motility other than that due to respiration-induced artifacts. No axons noticeably enlarged their tips, forming putative dystrophic endings. Following the first day of imaging, we imaged these axons repeatedly over several days to exclude the possibility that the immobility was due to phototoxicity (data not shown). These findings thus confirm that axons become rapidly immobilized, within a day or likely sooner, at the DREZ *in vivo*.

**Figure 2.**
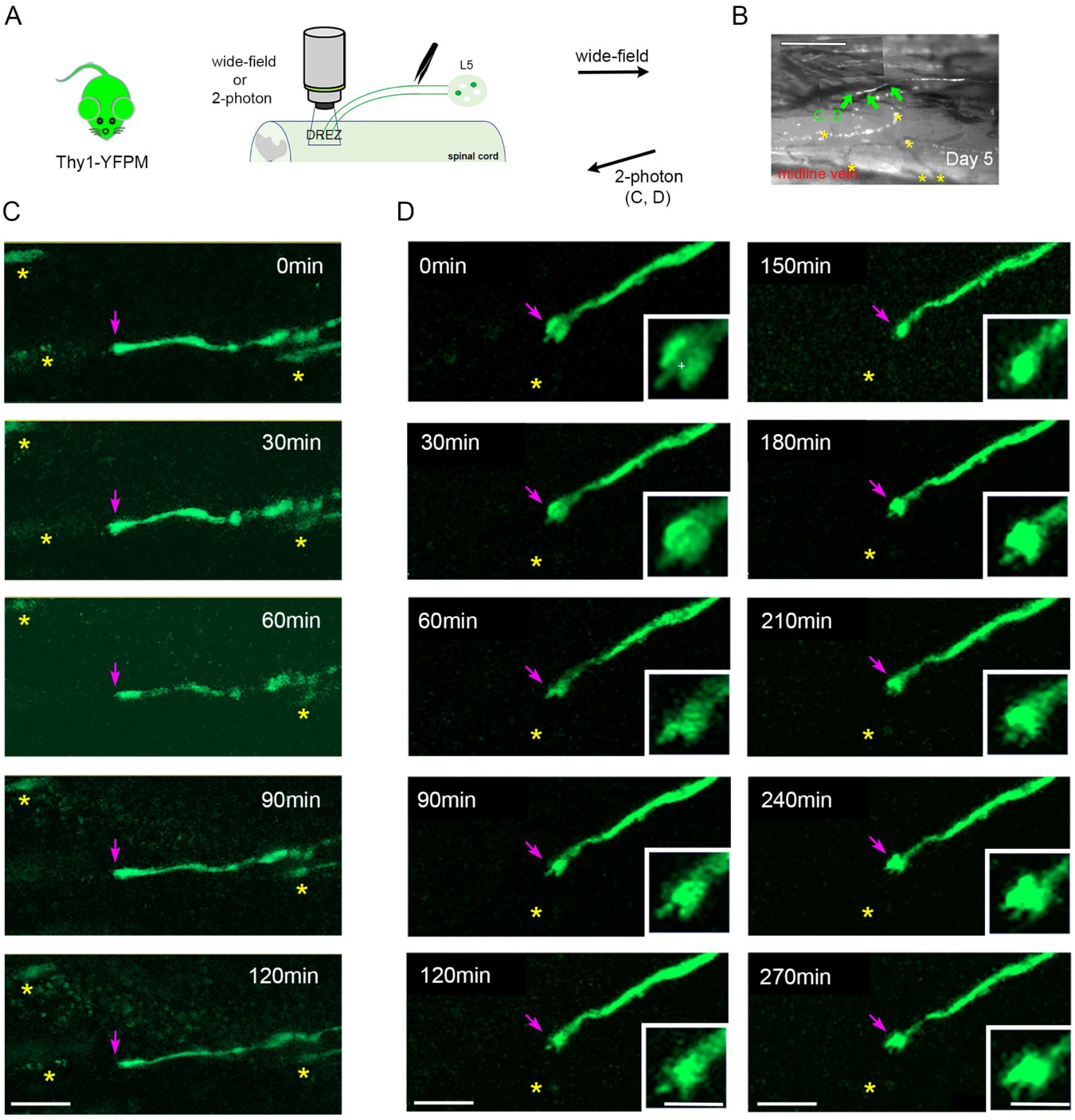
Two-photon imaging confirms rapid axon immobilization at the DREZ. **(A)** Schematic illustration of the experimental procedure for two-photon imaging of DR axons in live Thy1-YFPM mice after L5 root crush. **(B)** A widefield view of the DREZ area, taken at 5 dpi, showing an axon of interest (arrows) and fluorescent debris of degenerating old axons and a midline vein that served as landmarks (asterisks). Scale bar:100μm **(C, D)** Two-photon imaging of representative axon tips with either slender (C) or small roundish shape (D), taken every 30 min on 5 dpi. Arrows denote immobile axon tips and asterisks denote landmarks. Scale bar: 50μm (C, D), 20μm (D, insets)

### Relative positions of non-neuronal cells and arrested axons at the DREZ

To determine the extrinsic mechanism that rapidly immobilizes axons, we next examined various non-neuronal cells at the DREZ before and after DR crush. At 2 wpi, axons that stopped past the borderline were not associated with Schwann cell processes (Figure 3A, 3B), showing that they invaded CNS territory without leading, growth-supporting Schwann cells (Figure 3C; see also Figure 4A). These axons grew through the region of the DREZ, where astrocytes and oligodendrocytes were profuse, suggesting that these glia cannot rapidly immobilize axons (Figure 3A, 3B). Macrophages were far more numerous peripherally but notably sparse at axotomized DREZ (Figure 3B, Figure S1A), suggesting that growth-collapsing activity of macrophages ^21,22^ contributes minimally if at all to the axon arrest at the DREZ. Blood vessels, known for guiding axon growth ^23^, were also sparse along the root and at the DREZ (Figure S1D). Similarly, the two types of perivascular cells associated with blood vessels, pericytes and fibroblasts [i.e., vascular leptomeningeal cells (VLMCs)], were also scarce at both intact and axotomized DREZ (Figure S1B, S1D). These perivascular cells remained in association with the blood vessels after DR crush injury (Figure S1C), and we did not observe fibrotic scars filled with numerous macrophages, pericytes and fibroblast-like cells (Figure S1A, S1B), such as those elicited by spinal cord injury ^24^ or by severe root avulsion (Figure S1D). Because many axons are arrested at the DREZ, it is unlikely that non-neuronal cells that are sparse in the region can account for the rapid arrest of many axons (Figure 3C).

**Figure 3.**
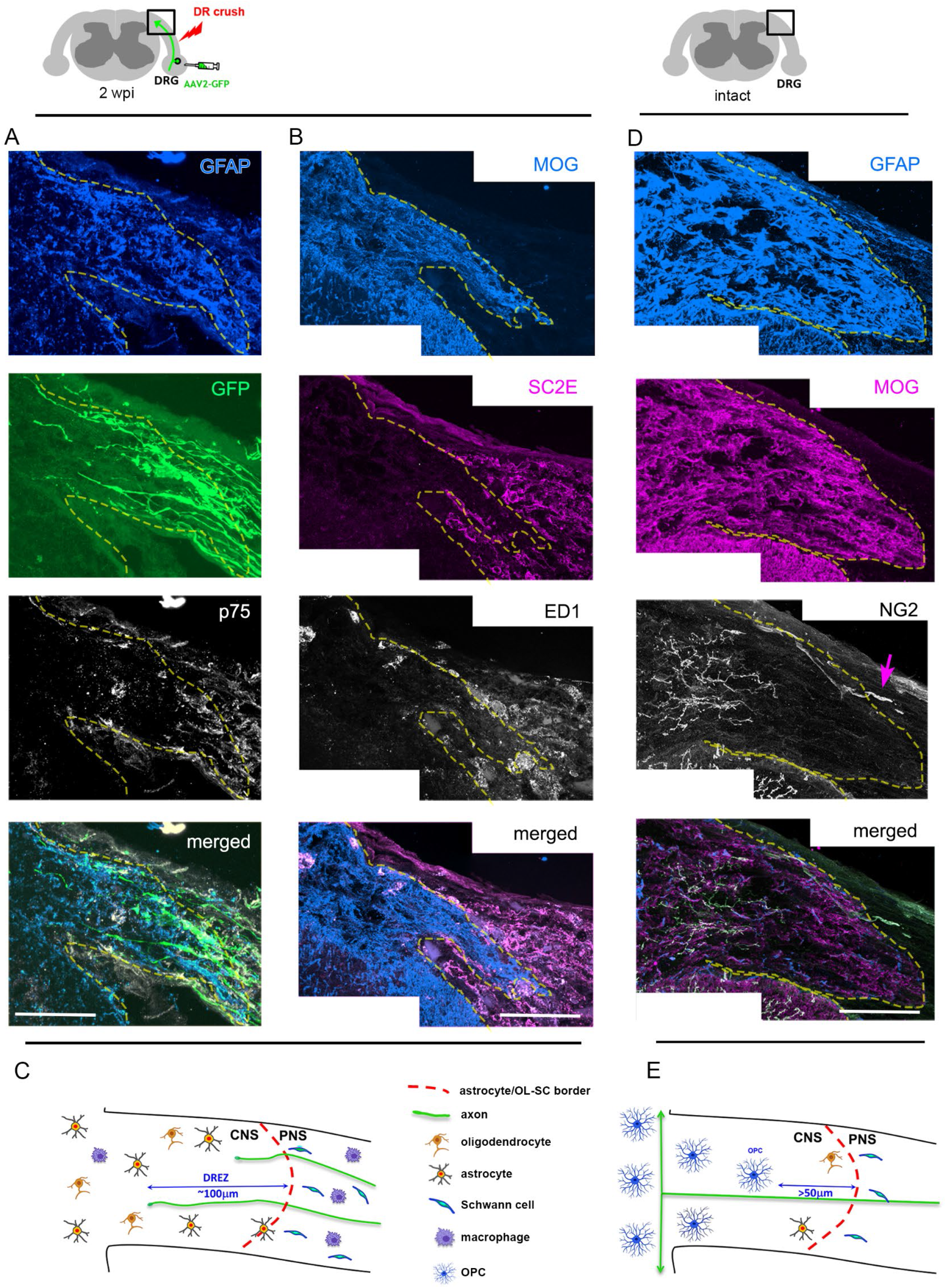
Relative positions of non-neuronal cells and arrested axons at the DREZ. **(A, B)** Transverse sections of a spinal cord 2 weeks after cervical root crush followed by intraganglionic injection of AAV2-GFP. **(A)** A DREZ area illustrating spatial relationships of arrested GFP+ axons (Green) with GFAP-labeled astrocytes (Blue) and p75-labeled Schwann cell processes (B/W). Many axons extend across the borderline (dotted line) by growing through astrocytes without association with leading Schwann cell processes. Scale bar: 50μm **(B)** A DREZ area illustrating spatial relationships of arrested axons (refer to A) with MOG-labeled oligodendrocytes (Blue), SC2E-labeled myelinating Schwann cells (Magenta), and ED1-labeled macrophages (B/W). Macrophages are sparse at the DREZ. Scale bar: 50μm **(C)** Diagram summarizing insignificant spatial relationships of arrested axons with oligodendrocytes, astrocytes, Schwann cells, and macrophages at axotomized DREZ. **(D)** A DREZ area of uninjured DRs illustrating that NG2-labeled OPCs (B/W) are absent at and immediately near the borderline (dotted line), denoted by astrocytes (Blue) and oligodendrocytes (Magenta), but abundant deeper at the DREZ (>50μm from the borderline). Arrow denotes NG2+ pericytes associated with a blood vessel. Scale bar: 50μm **(E)** Diagram illustrating the distance and position of OPC processes relative to the borderline (dotted line) at intact DREZ with no axon injury. See also Figure S1 and S2

**Figure 4.**
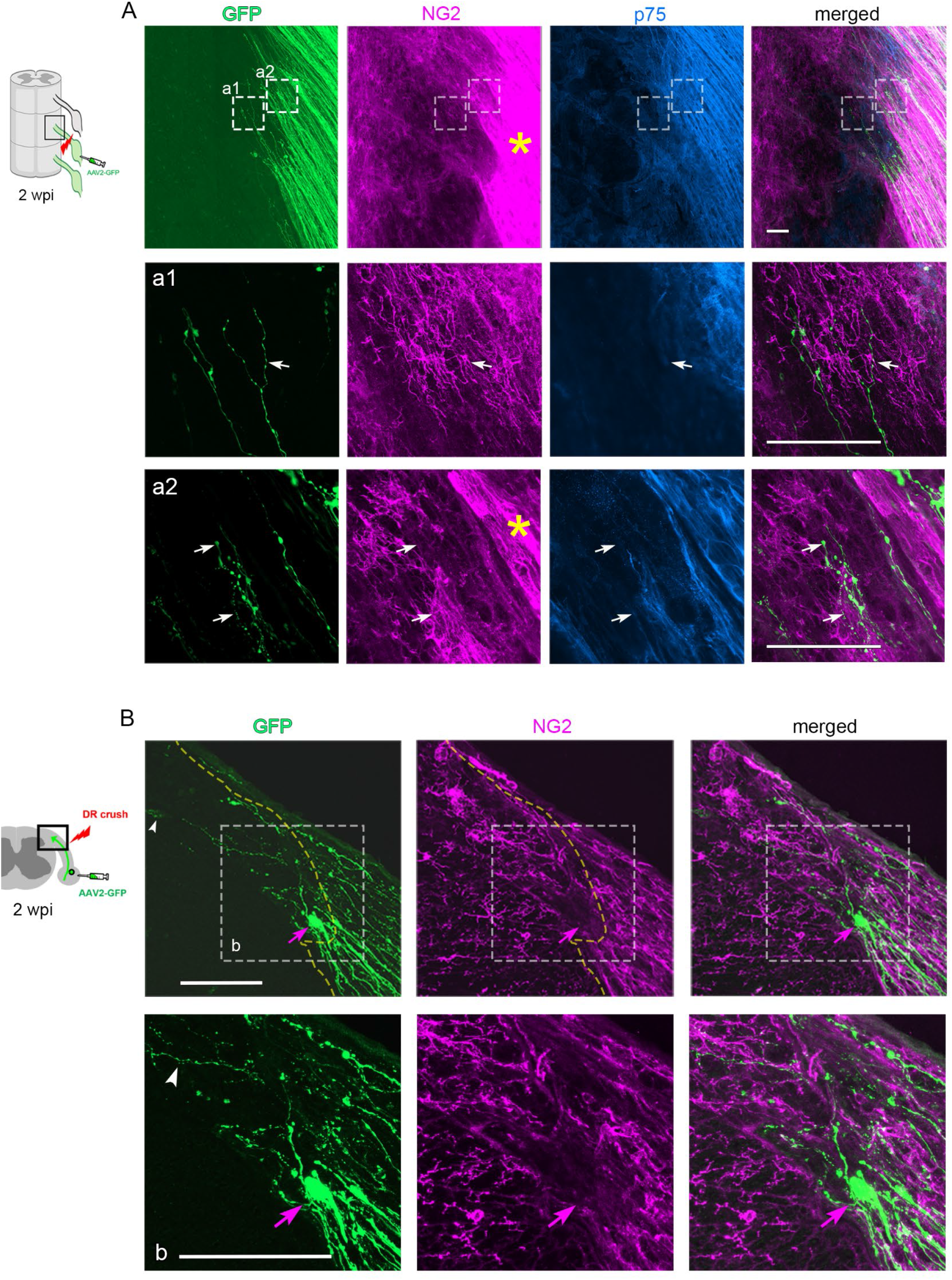
Stopped axons without dystrophic endings display extensive contacts with OPC processes. **(A)** Wholemount views of the DREZ area in low and high magnifications at 2 wpi, showing GFP+ axons (Green), NG2-labeled OPCs (Magenta), and p75-labeled Schwann cells (Blue). Arrows indicate examples of direct contacts between OPC processes and arrested axons. NG2 staining in the PNS (asterisks) is intense after DR injury but does not preclude identification of OPC processes. Scale bar: 100μm **(B)** Transverse sections in low and high (b) magnifications showing GFP+ axons (Green) and NG2-labeled OPCs (Magenta) at 2 wpi. Arrowheads indicate an example of many axons that stopped past the borderline (dotted line) in association with OPC processes. Arrows indicate a dystrophic ending formed at the borderline that, in contrast, lacks contacts with OPC processes. Scale bar: 50μm

Interestingly, NG2 staining showed that OPCs and their processes were absent at and within ∼50μm of the borderline, but abundant deeper in CNS territory (Figure 3D, 3E; Figure S2A-C). This distribution of resident OPCs before DR injury was consistent in all mice (n >20) and is noteworthy because many axons grow across the borderline and then stop deeper at the DREZ ^9^. OPCs are defined by their expression of both NG2 and PDGFRα, which are also expressed by other cell types ^24,25^. After DR injury, however, almost if not all macrophages were NG2-(Figure S1A). Both VLMCs and meningeal fibroblasts, which express PDGFRα but little if any NG2 ^26,27^, were immunostained weakly if at all by NG2 (Figure S1B). Schwann cells increased NG2 expression after injury, but they did not invade the DREZ (Figure 3). NG2+ pericytes, which do not express PDGFRα ^26^, were also sparse at the DREZ (Figure S1C). Furthermore, the perivascular configuration of pericytes was readily distinguished from the highly ramified morphology of OPCs (e.g., Figure 3D). Lastly, NG2 staining overlapped closely with PDGFRα staining at the DREZ (Figure S1B, S1C). Thus, almost all NG2+ cells at the DREZ were OPCs. Additionally considering that OPCs form synapses with axons, spatial analysis distinguishes OPCs from other non-neuronal cells.

### OPCs proliferate, migrate, and form a dense meshwork of processes after DR injury

Next, we examined responses of OPCs to DR injury in wholemounts, which permits comprehensive analysis of an entire root. We used Thy1-H mice in which subsets of degenerating or regenerating YFP+ axons are fluorescently labeled. In intact DRs, OPCs and their ramified processes were intraspinally distributed in a tile-like arrangement but were rare at the borderline (Figure S2A). At 2 wpi, many more OPCs formed a much denser meshwork of processes along the trajectory of degenerating distal axons (Figure S2B). OPC processes were now present at the borderline but remained relatively rare, compared to their density deeper in CNS territory. Also notable was an upregulated expression of NG2 proteoglycans in the PNS, presumably associated with denervated Schwann cells ^28^ and some fibroblasts ^29^. Schwann cell-associated NG2 staining did not preclude identification of OPC processes at the DREZ, however, because it was tube-like and distinctly brighter than that associated with OPCs, which was highly ramified and directionless (e.g., Figure 4A). Temporal analysis in transverse sections showed that by 5 dpi, when DR axons stop at the DREZ, OPC processes had migrated closer to the borderline, and by 2 wpi, their density deeper in CNS territory had increased (Figure S2C). OPCs also rapidly proliferated in response to the distant root injury; their proliferation peaked at 3 dpi, and their number doubled by 1 wpi (Figure S2D). These data suggest that most axons cross the borderline when OPCs are absent and then encounter OPCs.

### Stopped axons without dystrophic endings display extensive contacts with OPC processes

To determine if stopped axons are associated with OPCs, we labeled regenerating axons with an adeno-associated viral tracer, AAV2-GFP, and examined all detectable axon tips of entire roots in wholemounts at 2 wpi. Distal ends of almost all stopped axons were associated with a dense meshwork of OPC processes which made extensive contacts with axon endings and adjacent shafts (Figure 4A; n > 400 axons, 3 roots). Transverse sections also showed that all axons that crossed the borderline and stopped in CNS territory were in direct contact with OPC processes (Figure 4B). However, dystrophic endings at the borderline were not associated with OPCs (Figure 4B; n= >10). This finding is consistent with the absence of OPCs at the borderline and the lack of cellular contacts by dystrophic endings. These data suggest that axons occasionally stop by forming dystrophic endings embedded in collagen fibers at the borderline where OPCs are absent, whereas many others cross the borderline and then stop by making extensive contacts with OPCs.

### Stopped axons make synaptic contacts with OPC processes at the DREZ

Next, we used immunolabeling for synaptic markers to determine whether axon-OPC contacts are synaptic. Consistent with the ultrastructural observation of aggregated synaptic vesicles (Figure 1A), a synaptic vesicle marker, SV2, brightly labeled small swellings of axon tips and shafts tightly associated with OPC processes, when assessed in wholemounts at 2 wpi (Figure 5A, 5B). However, although less brightly, SV2 also labeled dystrophic endings and axon swellings that showed no obvious OPC contacts. Other vesicle markers, active zone and postsynaptic markers that we tested were also not convincingly selective (data not shown). Our attempts to trans-synaptically label OPCs by intra-DRG injections of pseudorabies or wheat-germ agglutinin (WGA)-expressing viral tracers were unsuccessful. Other investigators have also failed to provide light microscopic identification of neuron-OPC synapses, an anatomically and functionally weaker synapse than neuron-neuron synapses ^30,31^.

**Figure 5.**
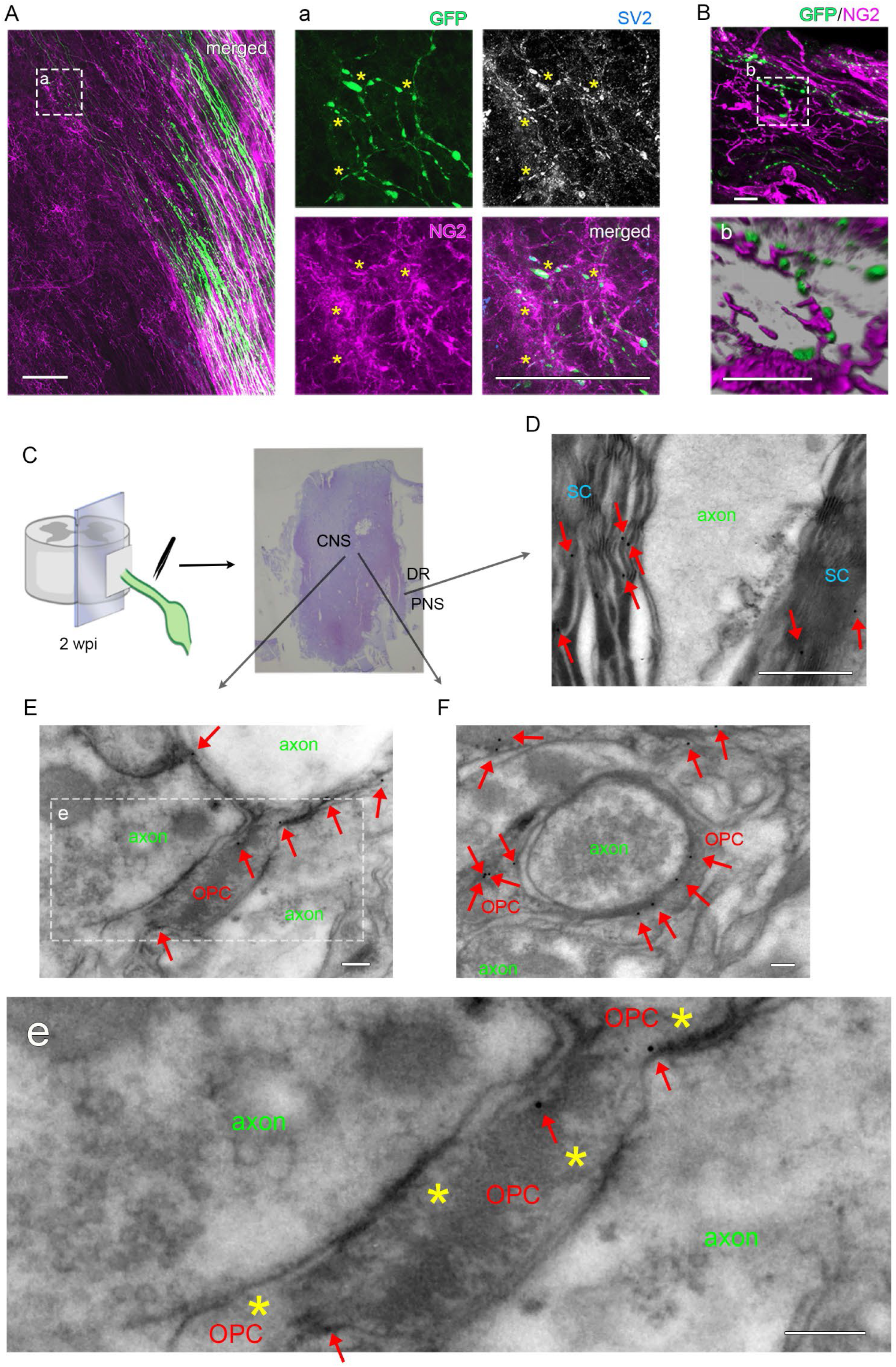
DR axons arrested at the DREZ form synapses with OPC processes. **(A)** Wholemount views of the DREZ area at 2 wpi in low and high magnifications (a), showing GFP-labeled axons (Green), SV2-labeled synaptic vesicles (B/W), and NG2-labeled OPCs (Magenta). Boxed area shows several axons with small swellings on axon tips and shafts intensely labeled for SV2 and tightly associated with OPC processes. Asterisks indicate same positions. Scale bar: 100μm (A), 50μm (a) **(B)** Volume rendered image (b) of a boxed area in B, illustrating direct contacts of axon swellings (Green) with OPC processes (Magenta). Scale bar: 5μm **(C)** Diagram showing procedures to prepare a longitudinal vibratome slice of dorsal spinal cord with attached DR at 2 wpi. **(D)** An electron micrograph of the PNS area showing gold particles (arrows) targeted to NG2 selectively labeling Schwann cells (SC). Scale bar: 500 nm **(E)** An area of the DREZ showing that OPC processes denoted by gold particles (arrows) form synaptic contacts with axons. Magnified view of boxed area (e) illustrates synaptic profiles (asterisks) with presynaptic clustering of synaptic vesicles and postsynaptic density. Scale bar: 100 nm **(F)** An electron micrograph illustrating varicosity-like synaptic profile formed by DR axon encircled by OPC processes identified by gold particles (arrows). Scale bar: 100 nm See also **Figure S3**

To obtain more convincing evidence, we performed immuno-EM. At 2 wpi, we prepared longitudinal slices of dorsal spinal cord containing a root and carefully examined the CNS and PNS areas across the borderline (Figure 5C). In the PNS, gold particles targeted to NG2 proteoglycans selectively labeled Schwann cells but not axons, confirming their specificity (Figure 5D). Remarkably, we frequently observed presynaptic profiles exhibiting vesicle clustering and a thin postsynaptic density in direct apposition to gold particles-labeled postsynaptic processes (Figure 5E). We also observed varicosity-like synaptic profiles, filled with numerous vesicles, formed on gold particles-labeled postsynaptic processes that encircled the axons (Figure 5F). Notably, these postsynaptic processes were thinner than axons, and their membranes, but not cytoplasm, were labeled by gold particles, consistent with the transmembrane location of NG2 proteoglycan ^32^. Because no other NG2+ cells form bona fide synapses, our immuno-EM identified OPCs as the postsynaptic cells that induce presynaptic differentiation of DR axons.

### OPCs form functional synapses with co-cultured DRG neurons

Next, we co-cultured OPCs and DRG neurons to determine whether they form functional connections. As DR axons become unable to penetrate the DREZ, ∼1 week after birth ^33,34^, we purified OPCs from spinal cords of p16-p18 mice and co-cultured them with purified DRG neurons from adult mice. >95% pure OPCs at 3 div were labeled specifically by both markers of OPCs, NG2 and PDGFRα. They resembled OPCs *in vivo* by exhibiting a non-overlapping distribution of bushy NG2+ processes (Figure S3A). A few dissociated neurons were then applied sparsely to the cultured OPCs and co-cultured for 5 days in proliferating medium. The DRG neurons extended relatively short axons on OPCs, which continued to proliferate. Of note, DRG axons frequently developed small swellings which were microtubule-deficient but SV2-enriched, a typical feature of presynaptic terminals (Figure S3B). We did not observe a large bulbous ending, reminiscent of dystrophic endings.

To determine whether these putative synaptic contacts were functional, we performed calcium imaging on the fifth day of co-culture. We loaded cultured cells with a fura-2 AM calcium indicator and locally infused 60mM KCL or 1mM glutamate. We observed, however, rapid calcium responses triggered not only from neurons but also from OPCs (data not shown). To specifically stimulate neurons, we applied capsaicin, which selectively depolarizes nociceptive neurons by activating TRPV1 receptors ^35^. Capsaicin did not elicit calcium responses in OPCs cultured without neurons (data not shown). Local capsaicin infusion elicited rapid calcium responses in neurons, followed by calcium responses in adjacent OPCs (Figure S3C). The responses in these OPCs were blocked by combined application of CNQX and AP5, antagonists of AMPA/kainate and NMDA receptors, consistent with earlier studies ^36,37^. Following washout of the antagonists, capsaicin-stimulation of the same neuron again evoked calcium responses in the same OPCs. These findings are consistent with an earlier study that demonstrated synaptic vesicle recycling and ultrastructural evidence of synaptic contacts in OPC-DRG cocultures ^38^. They indicate that OPCs form functional synapses with DRG neurons *in vitro* and reinforce the notion that axons may rapidly stop regenerating by forming synapses with OPCs.

### OPC ablation enables many axons to regenerate into the spinal cord

Because no OPC-associated synaptogenic inducer has been identified, we could not selectively inhibit synapse formation between OPCs and DR axons. As an alternative, we genetically ablated OPCs to determine whether they stop DR axons reentering spinal cord. OPC ablation *in vivo* has been difficult because they are highly proliferative and subject to tight spatial homeostasis: residual OPCs vigorously repopulate a depleted area ^39,40^. We therefore tried several transgenic strategies and diverse ablation paradigms. The best, yet incomplete, strategy was to use the PDGFRα-CreER driver and induce expression of diphtheria toxin subunit A (DTA). As DTA lacks the B subunit required for DT to penetrate membranes, ablation is restricted to cells synthesizing DTA proteins, which elicit apoptotic extirpation ^41^. We crossed PDGFRα-CreER and Rosa-DTA mice to generate PDGFRα-DTA mice and identified a 4-hydroxytamoxifen (4-HT) administration paradigm that elicited maximal ablation at 2 wpi (Figure 6A). Consistent with earlier studies that ablated OPCs by expressing DT receptors (DTR) or DTA ^42-44^, PDGFRα-DTA mice were grossly normal. NG2 and PDGFRα were markedly reduced as assessed by western blotting (Figure 6B) and immunostaining (Figure 6D, S4A). As anticipated, the extent of OPC ablation was quite variable, ranging between 20-80% at 2 wpi, as determined by mean fluorescence intensity of NG2 in the spinal cord. None of the administration regimens that we tried achieved more consistent or complete ablation. We therefore selected mice with at least 60% ablation at 2 wpi for further analysis (Figure 6C).

**Figure 6.**
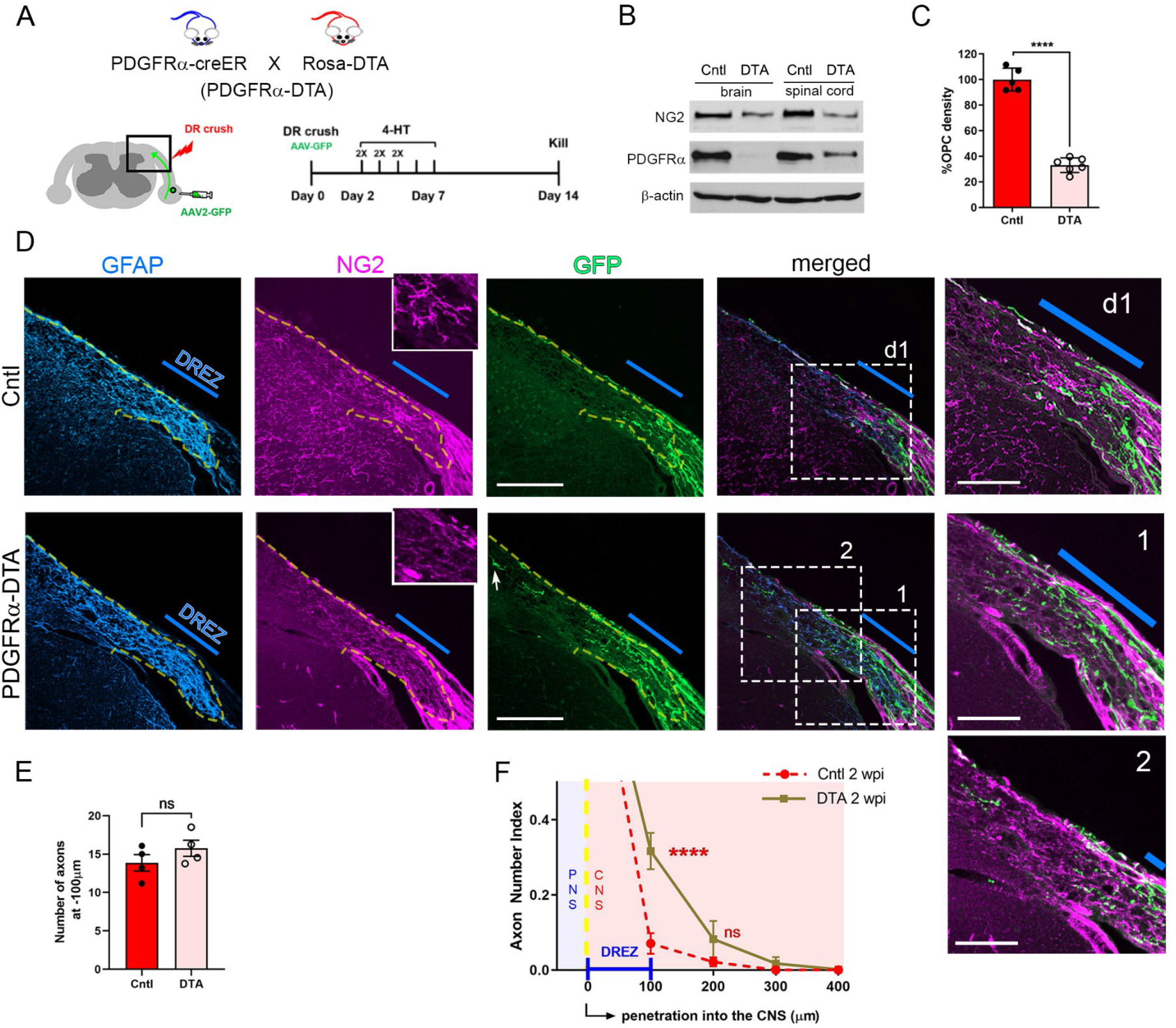
Genetic OPC ablation, although incomplete, markedly enhances regeneration across the DREZ. **(A)** Diagram illustrating experimental procedures to evaluate the effects of genetic OPC ablation at 2 weeks after C5-C8 root crush followed by AAV2-GFP injections to C5 and C7 DRGs. **(B)** Western blotting showing markedly reduced markers of OPCs, NG2, and PDGFRα, in PDGFRα-DTA mice. **(C)** Quantitative comparisons of OPC density measured by mean fluorescence intensity of NG2 staining in the spinal cord, illustrating ∼70% ablation of OPCs in PDGFRα-DTA mice used for the analysis. Cntl, 100.0±8.992; DTA, 33.07±5.725. Error bars represent SD. Unpaired t test (n=5 control, 6 DTA mice). ****p ≤ 0.00001 **(D)** Transverse sections of a control and a PDGFRα-DTA mouse with ∼80% OPC ablation at 2 wpi showing GFAP-labeled astrocytes (Blue), NG2-labeled OPCs (Magenta), and GFP+ axons (Green). Enlarged views of boxed area (d1, 1, 2) illustrate that more axons cross the border and DREZ in the PDGFRα-DTA mouse. Blue lines approximate ∼100μm wide region of the DREZ. Dotted lines denote borderline identified by GFAP staining. Insets are magnified views of the DREZ, illustrating residual and/or repopulating OPCs in PDGFRα-DTA mice. An arrow denotes axons extending unusually deeply into the dorsal funiculus. Scale bar: 100μm, 50μm (insets) **(E)** Quantification of GFP+ axons at ∼100μm proximal to the borderline in the PNS. Cntl, 13.86±1.075; DTA, 15.76±1.052. Error bars represent SEM (n = 4 mice/group). Unpaired Student’s t-test. ns, p > 0.05 **(F)** Quantitative comparisons of control and PDGFRα-DTA mice illustrating that OPC ablation triples the number of axons extending past the DREZ. Cntl: 0μm, 1.00±0; 100μm, 0.0765±0.02744; 200μm, 0.02128±0.01181; 300μm, 0.00±0.00. DTA: 0μm, 1.00±0.00; 100μm, 0.3167±0.0485; 200μm, 0.08238±0.04858; 300μm, 0.01723±0.01723; 400μm, 0.001375±0.001375. Error bars represent SEM (n=4 mice/group). Two-way repeated-measures ANOVA with Tukey’s multiple comparisons test. 100μm, ****p ≤ 0.00001; 200μm, ns, p > 0.05; 300μm See also **Figure S4, S5, S6**

At 2 wpi, some OPC processes usually remained at the DREZ even in mice with ∼80% ablation of OPCs (Figure 6D), presumably reflecting more vigorous proliferation of OPCs along the trajectory of degenerating injured axons. Remarkably, PDGFRα-DTA mice showed a greater number of axons at and beyond the DREZ than control mice (Figure 6D). The number of axons counted peripherally in PDGFRα-DTA mice did not differ from that in control mice, indicating that the enhanced regrowth was not due to an increase in axons regenerating peripherally (Figure 6E). Quantification of GFP+ axons showed that OPC ablation almost tripled the number of axons that extended past the DREZ (Figure 6F). Several considerations make the increase even more noteworthy. First, axons frequently extended unusually deeply, into the dorsal funiculus. Second, AAV2-transduced GFP+ axons are predominantly large-diameter axons whose growth capacity is particularly weak ^9^, yet even incomplete OPC ablation enabled ∼30% of these weak regenerators to cross the DREZ. Indeed, such a large increase rivaled the levels achieved by combining a conditioning lesion with simultaneous removal of myelin inhibitors and CSPGs ^9^.

### Enhanced regeneration is not due to indirect effects of OPC ablation

Recent scRNA sequencing studies indicated that PDGFRα is also expressed in VLMCs and that NG2 is also expressed in pericytes ^26,45,46^. This suggests that VLMCs and other PDGFRα+ fibroblasts are also deleted in PDGFRα-DTA mice. Additionally, NG2 staining will label OPCs and pericytes, whereas PDGFRα staining will label OPCs and VLMCs. Consistent with this notion, vascular NG2+/PDGFRα-labeling remains in PDGFRα-DTA mice, indicating that pericytes are not deleted (Figure S4A). PDGFRβ staining, which labels both pericytes and VLMCs, was reduced in PDGFRα-DTA mice, reflecting ablation of VLMCs (Figure S4B). In the PNS of uninjured control mice, we observed PDGFRα+ cells diffusely scattered between nerve fibers; these cells were not Schwann cells (Figure S4C), but were endoneurial fibroblasts (Figure S4D). PDGFα staining was markedly increased in the DR after root crush, reflecting rapid proliferation of endoneurial fibroblasts ^29^. We did not, however, observe a significant difference in root-associated PDGFRα staining between control and PDGFRα-DTA mice after injury (Figure S4D). Together, these data suggest that some PDGFRα+ fibroblasts are additionally deleted in PDGFRα-DTA mice. Their ablation, however, is highly unlikely to account for the enhanced regeneration, because VLMCs are sparse at intact and axotomized DREZ (Figure S1), and because axons regenerated normally along the roots in PDGFRα-DTA mice (Figure 6E).

Apoptosis does not induce an inflammatory response ^47,48^. In accordance, and consistent with earlier studies of OPC ablation ^42,49,50^, the inflammatory responses of PDGFRα-DTA mice were comparable to those of control mice: the morphology, distribution, and numbers of microglia and macrophages in the spinal cord and DRs were not significantly different (Figure S5A-B, S5D-F). Astrocytes, which inhibit Schwann cell migration ^51^, also showed no differences in morphology, density, or reactivity (Figure S5C, S5G). NG2 staining in the PNS of PDGFRα-DTA mice was slightly diminished, as compared to control mice, at 2 wpi (Figure S6A, S6E). Because Schwann cells do not express PDGFRα mRNAs ^52^, this reduction likely reflects ablation of some PDGFRα+ endoneurial fibroblasts. Consistent with this notion and Figure S4C, the density and reactivity of Schwann cells in DTA and control mice were not significantly different (Figure S6B-C, S6F-G). Lastly, NG2, also known as CSPG4, is a constituent of growth inhibiting CSPGs ^2,53^. It is conceivable, therefore, that a reduction of NG2/CSPG4 is the basis of enhanced regeneration by OPC deletion. We found, however, abundant expression of CSPGs in PDGFRα-DTA mice, as indicated by CS56 immunoreactivity (Figure S6D, S6H). Moreover, we were unable to achieve more robust regeneration even after broad removal of all CSPGs ^9^. Collectively, these data exclude that the enhanced regeneration was due to indirect effects on axons caused by ablating OPCs.

### Axons continue regenerating intraspinally through areas lacking OPCs

Stopped axons remained remarkably stable at the DREZ for at least 4 months after injury ^6^. We also reported that axons remain immobilized for at least 4 weeks at the DREZ, despite simultaneous and prolonged removal of myelin inhibitors and CSPGs ^9^. If OPCs play a primary role in stopping axons, then axons may continue to grow with prolonged ablation of OPCs, even in the presence of other extrinsic inhibitors. To test this notion, we examined PDGFRα-DTA mice after longer survival, 4 wpi (Figure 7A). As anticipated, OPCs widely repopulated and re-formed an OPC meshwork in most areas of the spinal cord; their overall densities corresponded to ∼70% of those in control mice (Figure 7B). Of note, however, OPC repopulation was not uniform; there were areas in which OPCs remained reduced or sparse (Figure 7D, S7A-B).

**Figure 7.**
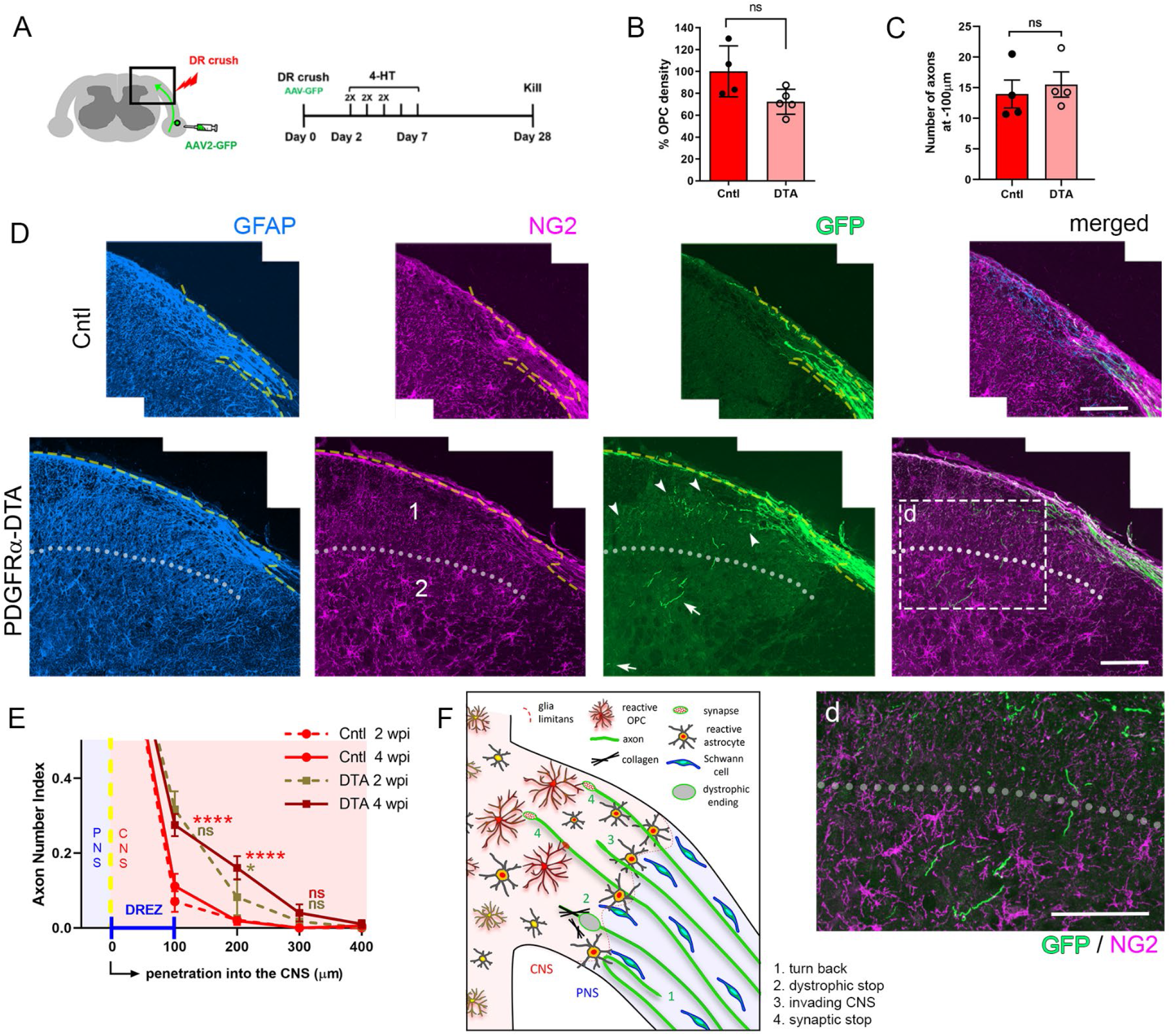
Axons continue regenerating intraspinally through areas lacking OPCs. **(A)** Diagram illustrating experimental procedures to evaluate the effects of genetic OPC ablation at 4 weeks after C5-C8 root crush followed by AAV2-GFP injections into C5 and C7 DRGs. **(B)** Quantitative comparison of OPC density, measured by mean fluorescence intensity of NG2 staining in the spinal cord, illustrating ∼30% ablation of OPCs in PDGFRα-DTA mice used for the analysis. Cntl, 100.0±23.35; DTA 72.31±11.4. Error bars represent SD. Unpaired t test (n=4 control, 5 DTA mice). ns, p > 0.05 **(C)** Quantification of GFP+ axons at ∼100μm proximal to the borderline, illustrating insignificant difference between control and PDGFRα-DTA mice. Cntl, 13.98±2.281; DTA 15.53±2.063. Error bars represent SEM (n = 4 mice/group). Unpaired Student’s t-test. ns, p > 0.05 **(D)** Transverse sections from a control and a PDGFRα-DTA mouse, illustrating massive repopulation of OPCs and GFP+ axons that regenerate remarkably deeply into the dorsal horn (arrows). There are noticeably fewer OPC processes where more axons (arrowheads) regrow within the spinal cord (area 1) than where fewer axons (arrows) regenerate (area 2). Enlarged view of boxed area (d) illustrates regenerated GFP+ axons in association with repopulated OPC meshwork. Scale bar: 100μm **(E)** Quantitative comparison of control and PDGFRα-DTA mice illustrating that ∼20% axons continue to regenerate past the DREZ in DTA mice during the additional two weeks. Error bars represent SEM (n=4 mice/group). Two-way ANOVA with Sidak’s multiple comparisons test. 100μm; ns, p > 0.05 (DTA 4 wpi vs. DTA 2 wpi); ****p ≤ 0.00001 (DTA 4 wpi vs. control 4 wpi), 200μm; ****p ≤ 0.00001 (DTA 4 wpi vs. control 4 wpi), *p ≤ 0.01 (DTA 4 wpi vs. DTA 2 wpi), 300μm; ns, p > 0.05 (DTA 4 wpi vs. control 4 wpi, DTA 2 wpi). **(F)** Schematic illustration of the proposed model. See also Figure S7.

Consistent with our previous study, in control mice, regeneration across the DREZ did not increase between 2 wpi and 4 wpi (Figure 7D, 7E). In striking contrast, in PDGFRα-DTA mice, GFP+ axons more frequently extended significantly deeper into the dorsal funiculus at 4 wpi than at 2 wpi (Figure 7D, S7A-B) and occasional axons regrew remarkably deeply into the dorsal horn (Figure 7D). Intraspinal growth of DR axons that reached deep layers of the dorsal horn was indeed extraordinary, because it was not achieved even by a nerve conditioning lesion ^9^ or PTEN deletion (Kim et al, in preparation). Also notable was a conspicuous reduction of OPC processes in areas where more axons grew deeper (Figure 7D). Furthermore, axons that extended through OPC-deficient areas terminated in OPC-rich regions (Figure S7A-B). The number of axons extended from the PNS did not increase (Figure 7C) nor did the number of axons counted at 100μm from the borderline in the CNS (Figure 7E). Therefore, the enhanced regeneration was due to sustained growth of the axons already present at the DREZ at 2 wpi, rather than by axons that freshly crossed the borderline. Taken together, these observations suggest that some axons could continue regenerating into the area of spinal cord where residual or repopulating OPCs were relatively rare.

## DISCUSSION

Here, we identify OPCs as the postsynaptic cells arresting sensory axons regenerating into the spinal cord. Our studies provide additional insights into the regeneration failure at the DREZ, prompting us to propose a new model (Figure 7F): Regenerating axons exiting basal lamina tubes at the borderline turn around or stop immediately by forming dystrophic endings if their growth is obstructed by collagen fibers. Most axons cross the borderline into CNS territory, albeit extending slowly due to poor regenerative ability, extrinsic growth-inhibitory molecules and lack of leading Schwann cells. These axons stop as they encounter a dense meshwork of OPC processes that induce presynaptic differentiation. Concurrent induction at multiple axon-OPC contacts evokes rapid and persistent synaptic stabilization.

Our findings definitively demonstrate that DR axons stop in two distinct ways, forming dystrophic or synaptic contacts. Our EM studies revealed numerous synapses or synapse-like profiles, but no dystrophic endings, in the CNS beyond the borderline. Other investigators also observed synapse-like profiles (termed synaptoids) and postulated the presence of “physiological stop signals” arresting DR axons ^54,55^. Together, we suggest that dystrophic endings form exclusively at the border area where OPCs are absent, but collagen fibers are abundant. Considering the much slower axon degeneration and weaker inflammatory responses in the CNS than PNS ^56^, the extent of intraneural fibrosis is likely high at the border and peripherally along the root ^56^. Accordingly, endoneurial fibroblasts proliferate rapidly after peripheral nerve injury ^29^, and we observed a marked increase of PDGFRα+ fibroblasts peripherally after DR injury. Moreover, our ultrastructural surveys frequently revealed collagen fibers peripherally but rarely at the DREZ. One speculative scenario is, therefore, that regenerating axons exiting basal lamina tubes at the borderline are occasionally obstructed by collagen fibers. Interestingly, extensive fibrosis inhibits OPC migration ^57^, suggesting that collagen fibers may trap dystrophic endings and also repel OPCs from the PNS.

Approximately 70% axons did not turn around or stop at the borderline, but invaded CNS territory. This is surprising considering the limited ability of axons to cross astrocyte-Schwann cell interfaces ^58,59^. However, axons can use bridges formed by astrocytes to cross the DREZ ^60^ and spinal cord lesions ^61^. Astrocytic processes invade peripherally after DR injury and are frequent within Schwann cell basal lamina tubes ^62^. We suggest, therefore, that astrocytic guidance is initiated peripherally and that it enables many axons to cross the borderline.

When axons crossed the borderline (∼5 dpi), proliferating OPCs filled most regions of the DREZ with a dense meshwork of highly ramified processes. Axons that invaded CNS territory would, therefore, almost immediately encounter OPCs and make multiple contacts with OPC processes. If, as we think likely, each axon-OPC contact elicits synaptic induction, then exuberant synaptogenic signals will easily overpower weak growth-promoting stimuli in CNS territory, thereby rapidly arresting axons. OPC-mediated synaptic arrest can, therefore, explain how axons are rapidly immobilized and how arrested axons can be maintained for a remarkably long time without degenerating. Supporting this notion are reports that presynaptic differentiation is rapidly induced within one hour after target contact ^63,64^. OPC synapses are formed on multiple discrete sites along unmyelinated axons, where vesicular glutamate release can elicit local Ca signaling on OPCs ^37,65^. These neuron-OPC synapses are maintained and transferred to the daughter OPC cells when OPCs divide ^66,67^.

OPC-mediated synaptic arrest is also consistent with recent studies implicating synaptic suppression as an important determinant of axon regeneration. DRG neurons downregulate presynaptic proteins essential for synaptic transmission, such as α2δ calcium channel subunits, as they regain the ability to grow their axons ^68^. Genetic or pharmacological targeting of α2δ or another active zone protein, Munc13, promotes regeneration of dorsal column axons after spinal cord injury ^68,69^. Intact axon branches maintaining synaptic connections similarly suppress regeneration of injured axon branches ^70^.

We suggest that OPCs stop regenerating axons, and that they do so by inducing synaptic differentiation. Silver’s group reported the seemingly analogous finding that spinal cord axons dying back after injury stop degenerating by making synapse-like contacts with OPCs ^38^. NG2/CSPG4 appears to mediate this process, termed synaptic entrapment, because knockout mice globally lacking NG2/CSPG4 show reduced entrapment, lengthened dying-back, and more robust intraspinal regeneration ^38^. Notably, however, synapses with clear ultrastructural features were not observed (Jerry Silver, personal communication). OPC-neuron synapses form readily in the absence of NG2/CSPG4 ^71^. Additionally, taking into account that the study from the Silver group examined degenerating, not regenerating, axons, we posit that NG2/CSPG4-mediated synaptic entrapment is not identical to OPC-mediated synaptic arrest. It remains to be determined if intraspinal axons stop regenerating by forming synapses with OPCs.

How OPCs induce presynaptic differentiation remains speculative. OPCs express several synaptogenic molecules, including Nlgn1-3 (Neuroligin1-3), Cadm1a/1b (SynCAM1a/1b), Lrrc4 (NGL-3), and Lrrtm 1/2 ^72^. Neuroligin and SynCAM are transsynaptic adhesion molecules that trigger presynaptic assembly when expressed in non-neuronal cells ^73,74^. Also undetermined is whether the formation of OPC-DR axon synapses requires synaptic transmission or activity. Notably, inhibition of synaptic transmission promotes regeneration of DC axons ^68,69^, whereas chemogenetic activation of DRG neurons combined with CSPG removal enhances regeneration across the DREZ ^75^. Although neuron-neuron synaptogenesis proceeds in the absence of neuronal activity ^76,77^, neuronal activity levels may have a differential effect on synapse number and maturation in different contexts ^78^. It will, therefore, be interesting to determine how activity levels of DRG neurons affect OPC-mediated synaptic arrest.

To verify the dominant role of OPCs, we sought to determine whether a nerve conditioning lesion would enhance regeneration if OPCs are depleted. However, because a conditioning lesion reduced the efficiency of OPC deletion, we could not obtain conclusive results (data not shown). Attempts to survey the mobility of axons arriving at the DREZ were also unsuccessful, because too many OPCs remained at 5 dpi. It is worth emphasizing, therefore, that we observed enhanced regeneration despite the continued presence of ∼30% OPCs at 2 wpi and ∼70% OPCs at 4 wpi. Lastly, the ability of OPCs to differentiate into myelinating oligodendrocytes is limited, although surplus numbers are rapidly generated after demyelination ^79^. The function of OPC-neuron synapses also remains enigmatic ^36^. Future efforts to determine if OPCs elsewhere in the CNS curtail axon plasticity and/or regeneration, may have significant basic science and clinical implications.

### Limitations of the study

EM remains the only unbiased method to unequivocally identify synaptic structures and partner cells, but ultrastructurally identifying a cell type is challenging. Our immunoEM identified axon contacts with NG2+ cells exhibiting apparent synaptic features. Although DRG neurons and OPCs were not specifically labeled in the study, it is highly likely that these synaptic contacts are made by regenerating DR axons and OPCs because no other axons are present at the axotomized DREZ and no other NG2+ cells form bona fide synapses. Nevertheless, it is necessary to confirm their identity unambiguously. In addition, although our light microscopic, in vitro, and EM studies all suggest an extensive incidence of axon-OPC synapses, we could not confirm this notion with immunoEM. Advanced technologies, such as genetic labeling, correlative light and electron microscopy and volume imaging will be able to demonstrate definitively the location, incidence, and cellular identity of axon-OPC synapses. Further limitations of the present study include incomplete OPC ablation and concomitant ablation of PDGFRα+ fibroblasts. Future studies that completely and specifically ablate OPCs will confirm and evaluate more fully the effects of OPC ablation on axon regeneration.

## ACKNOWLEDGMENTS

We thank Drs. Alan Tessler, Tim Himes, George Smith, and Gareth Thomas for critical reading of the manuscript. We thank Dr. Dwight Bergels for guinea pig anti-NG2 antibody, Drs. Dewight Williams and Biao Zuo at the Penn Electron Microscopy Resource Laboratory for technical support for TEM and immunoEM. We sincerely thank Dr. Ahmet Hoke for motivating the study, Drs. Jerry Silver and Theresa Connors for earlier immunoEM efforts, Dr. Michael Lane for pseudorabies virus, Dr. Jian Zhong for WGA-transsynaptic virus and Dr. George Smith for scAAV2-GFP. This work was supported by grants from NIH R01 (NS079631) and Shriners Children’s to Y-J Son. H.K. was a recipient of Craig H Neilson postdoctoral fellowship (280274).

## AUTHOR CONTRIBUTIONS

Conceptualization, Y-J.S.; methodology, S.K., H.H.; investigation, H.K., A.S., J.X., S.H., J.Z., Y-J.S.; writing—original draft, H.K.; writing—review and editing, Y-J.S.; funding acquisition, Y-J.S., H.K.; resources, S.K.,; supervision, Y-J.S.

## SUPPLEMENT FIGURE LEGENDS

**Figure S1.**
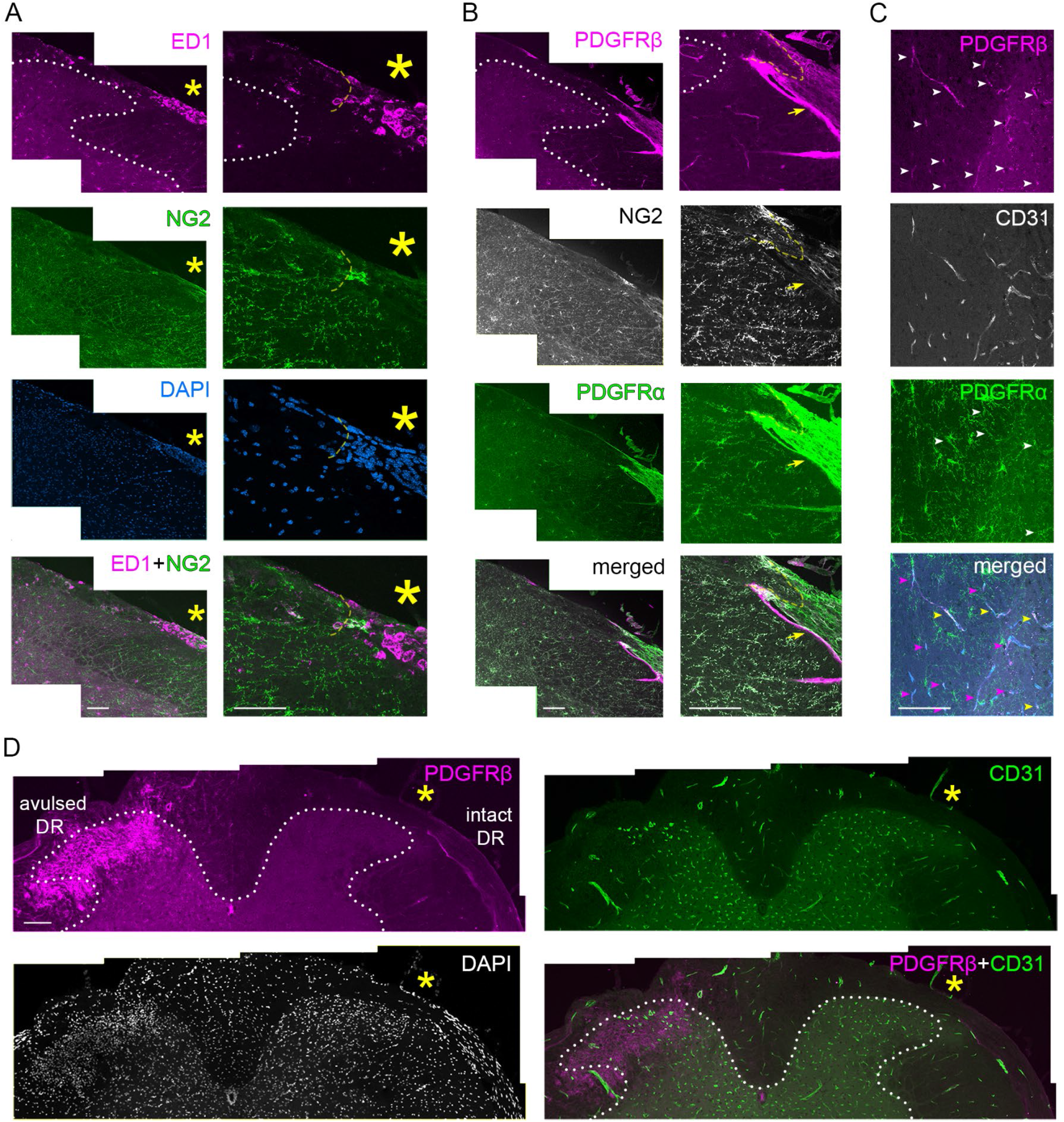
Macrophages, pericytes, fibroblast-like cells, and blood vessels are sparse at intact and axotomized DREZ. **(A)** Transverse sections of a spinal cord at 2 wpi, showing low and high magnification views of DREZ area. Asterisks denote numerous macrophages (Magenta) enriched peripherally along the root. Macrophages are sparse at the DREZ and NG2+ (Green). Yellow dotted line denotes borderline identified by peripherally enriched nuclei. White dotted line denotes grey matter. Scale bar: 100μm **(B)** Transverse sections of a spinal cord at 2 wpi, showing low and high magnification views of the DREZ area. Pericytes and VLMCs, labeled by PDGFRβ (Magenta), are sparse at the DREZ and appear to be perivascular as indicated by those within the spinal cord. Arrows denote PDGFRβ+/NG2-/PDGFRα+ meningeal fibroblasts associated with the nerve root. NG2 and PDGFRα staining overlaps closely at the DREZ. Because few if any other types of NG2+/PDGFRα+ cells (i.e., VLMCs) are present at the DREZ, almost all NG2+ cells at the DREZ are OPCs and their processes. Scale bar: 100μm **(C)** Transverse sections of a spinal cord at 2 wpi, showing high magnification views of grey matter. All pericytes and VLMCs, labeled by PDGFRβ (White arrowheads), remain associated with blood vessels (B/W) after DR injury. Some PDGFRβ labeling is not co-localized with PDGFRα staining (Magenta arrowheads), consistent with the notion that VLMCs, but not pericytes, express PDGFRα. Scale bar: 100μm **(D)** Low magnification views of transverse sections of a spinal cord showing an intact (right) and severely avulsed (left) root. Glial/fibrotic scar was intentionally elicited by severely avulsing C5-C6 roots to damage dorsal spinal cord. The scar is filled with numerous PDGFRβ+ cells. In contrast, PDGFRβ+ cells are sparse at the DREZ (asterisks) and spinal cord with an intact root or after root crush (**B**). Note also that blood vessels (Green) are sparse at the DREZ. Scale bar: 100μm

**Figure S2.**
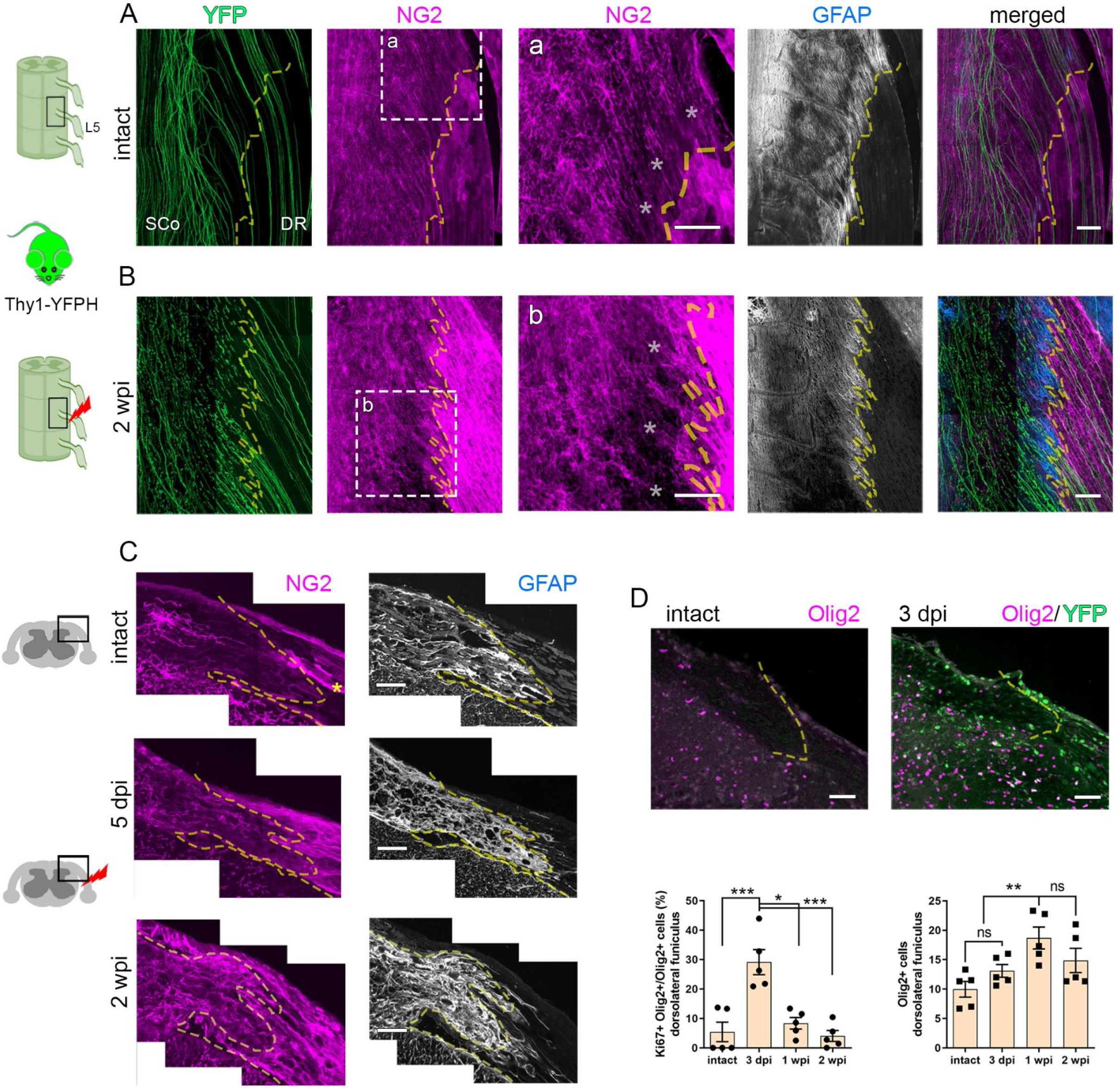
OPCs proliferate, migrate, and form a dense meshwork of processes after DR injury. **(A, B)** Wholemount views of intact (A) and axotomized (B) DREZ area showing subsets of YFP+ axons (Green) and the distribution of NG2-labeled OPCs and their processes (Magenta). Dotted line approximates the borderline denoted by GFAP-labeled astrocytes (B/W, Blue). Enlarged images (a, b) show area near the borderline (asterisks) and deeper in CNS territory. Scale bar: 100μm **(C)** Transverse sections of intact or axotomized DREZ at 5 dpi and 2 wpi, co-labeled for NG2 (Magenta) and GFAP (B/W, Blue), illustrating temporal development of a dense meshwork of OPC processes populating the DREZ after DR injury. Asterisk denotes NG2-labeled blood vessels. Scale bar: 50μm **(D)** Quantification of OPC proliferation by counting nuclei co-labeled by Olig2, a marker of OPCs and oligodendrocytes, and Ki67, a marker of cell proliferation. N= >3 sections/each mouse, 5 mice/each time point. Ki67+Olig2+: intact, 5.405±3.310; 3 dpi, 29.23±4.260; 1 wpi, 8.359±1.945; 2 wpi. 4.047±1.845. Olig2+: intact, 9.967±1.327; 3 dpi, 13.09±1.082; 1 wpi, 18.68±1.859; 2 wpi 14.87±2.065. Error bars represent SEM. p values are calculated by one-way repeated measures ANOVA with Tukey’s post hoc test. ns, p > 0.05; *p ≤ 0.01, **p ≤ 0.001, ***p ≤ 0.0001. **(D)** Quantification of OPC proliferation by counting nuclei colabeled by Olig2, a marker of OPCs and oligodendrocytes (D1), and Ki67, a marker of cell proliferation (D2), illustrating rapid proliferation of OPCs after DR injury that peaks at 3 dpi (D3) and doubles the number of Olig2+ cells by 1 wpi (D4). N= >3 sections/each mouse, 5 mice/each time point. Error bars represent SEM. p values are calculated by one-way repeated measures ANOVA with Tukey’s post hoc test. ns, p > 0.05; *p ≤ 0.01, **p ≤ 0.001, ***p ≤ 0.0001

**Figure S3.**
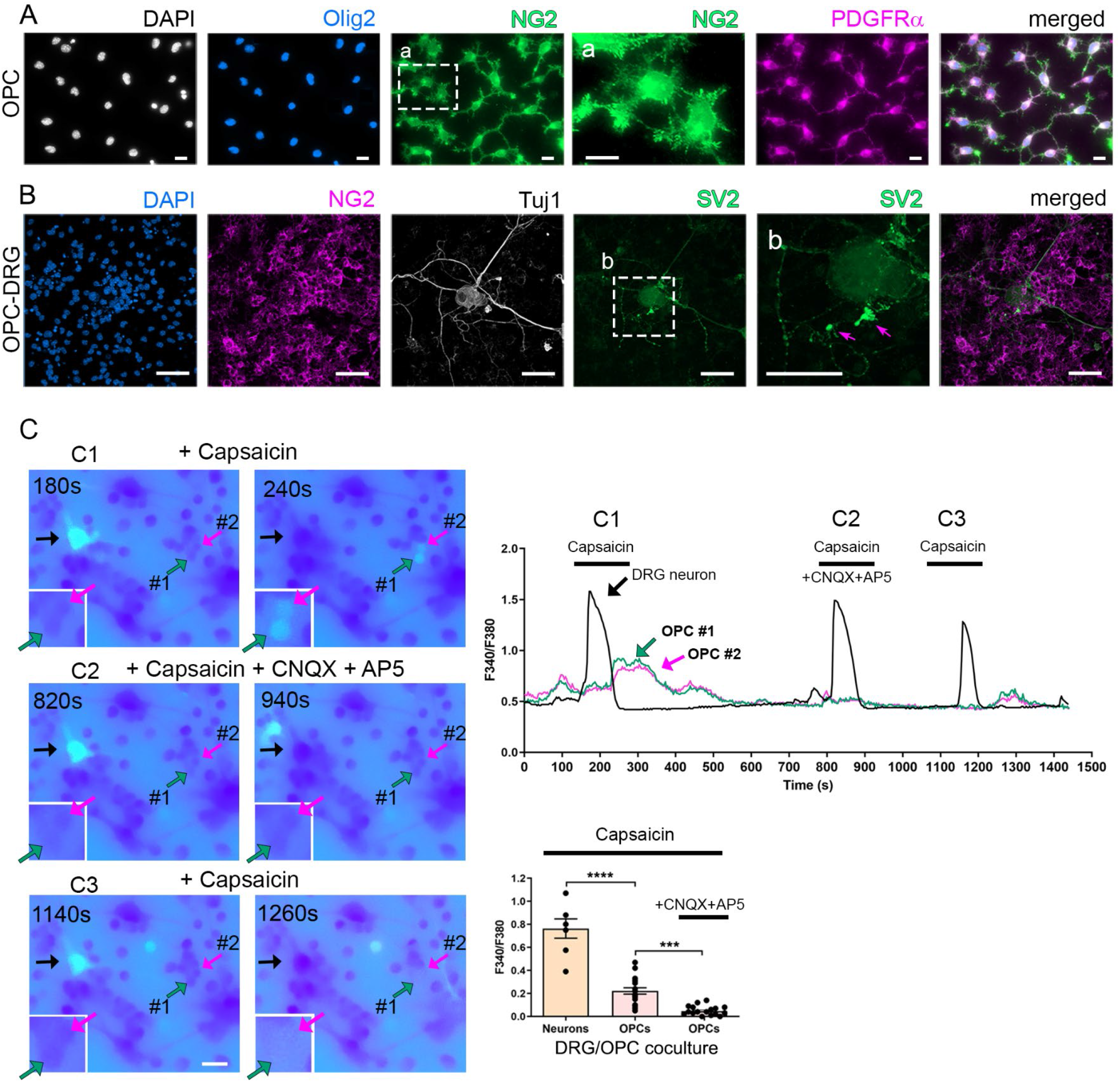
OPCs form functional synapses with co-cultured DRG neurons. **(A)** >95% pure OPCs, cultured from spinal cords of p16-18 mice, immunolabeled at 3 div for DAPI, Olig2, NG2 and PDGFRα. Enlarged view (a) of boxed area illustrates OPCs with bushy processes resembling OPCs *in vivo*. Scale bar: 20μm **(B)** Purified OPCs and DRG neurons co-cultured for 5 days, showing DAPI, NG2-labeled OPCs, Tuj1-labeled axons, and SV2-labeled synaptic vesicles. Magnified view (b) of boxed area illustrates microtubule-deficient but SV2-enriched axonal swellings (arrows) formed on OPCs. Scale bar: 50μm **(C)** Representative calcium imaging recordings of OPC-DRG neurons co-cultured for 5 days, illustrating temporal calcium responses of a neuron (Black arrows) and two adjacent OPCs (Green and Magenta arrows), following capsaicin stimulation of neurons without (C1) or with antagonists of AMPA/kainate and NMDA receptors (C2), CNQX and AP5. Following washout of antagonists (C3), capsaicin stimulation of the same neuron again elicits calcium responses in the same OPCs. Graph shows comparisons of capsaicin-elicited calcium responses in DRG neurons and OPCs. n = 7-18. Neurons, 0.76±0.05; OPCs, 0.22±0.06; OPCs, 0.05± 0.04. Error bars represent SEM. p values are calculated by one-way repeated measures ANOVA with Tukey’s post hoc test. ***p ≤ 0.001, ****p ≤ 0.0001. Scale bar: 30μm

**Figure S4.**
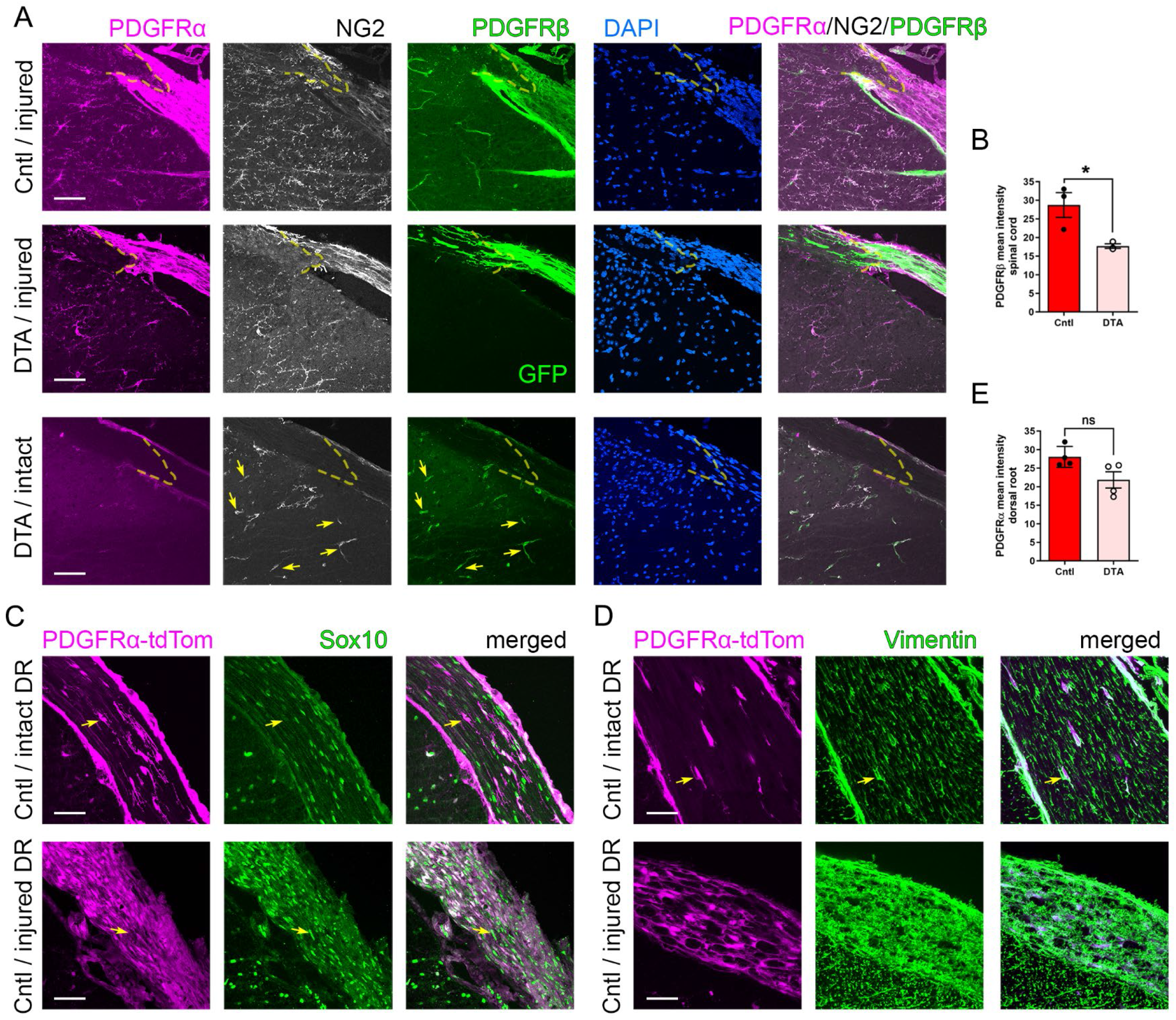
Pericytes and ablation of PGFRα+ cells in PDGFRα-DTA mice. **(A)** Representative high magnification views of the DREZ area in a control and two PDGFRα-DTA mice, co-labeled for PDGFRα, NG2, and PDGFRβ. A PDGFRα-DTA mouse with ∼40% OPC ablation at 2 wpi shows PDGFRα staining closely overlapped with NG2 staining, illustrating ablation of PDGFRα+ cells. Bottom panels show the DREZ area of an uninjured PDGFRα-DTA mouse with ∼80% OPC ablation, illustrating unaffected pericytes. NG2+/PDGFRβ+ cells with perivascular morphology (i.e., pericytes; arrows) remain intact, despite almost complete ablation of PDGFRα+ cells in the DREZ area of the DTA mouse. Scale bar: 50μm **(B)** Quantitative comparisons of control and PDGFRβ-DTA mice, illustrating a reduction of PDGFRβ immunoreactivity in the spinal cord of DTA mice. Quantification of the DREZ area is not provided due to paucity of PDGFRβ+ cells at the DREZ in both control and DTA mice. Cntl, 28.74±3.344; DTA, 17.69±0.6118. Unpaired Student t-test (n=3 control, 3 DTA mice). Error bars represent SEM. *p ≤ 0.01. **(C)** High magnification views of dorsal roots of a PDGFRα-tdTomato mouse, showing that PDGFRα+ cells are sparse in an intact dorsal root but abundant in a crushed root at 2 wpi. PDGFRα+ cells are not labeled by Sox10 (arrows), indicating that they are not Schwann cells. Scale bar: 50μm **(D)** High magnification views of intact and crushed dorsal roots of a PDGFRα-tdTomato mouse, showing that PDGFRα+ cells are co-labeled by a fibroblast marker, vimentin (arrows). Scale bar: 50μm **(E)** Quantitative comparisons of control and PDGFRα-DTA mice, illustrating insignificant difference in PDGFRα immunoreactivity in the dorsal root. Cntl, 28.04±1.406; DTA. 21.83±2.197. Unpaired Student t-test (n=4 mice/group). Error bars represent SEM. ns, p > 0.05

**Figure S5.**
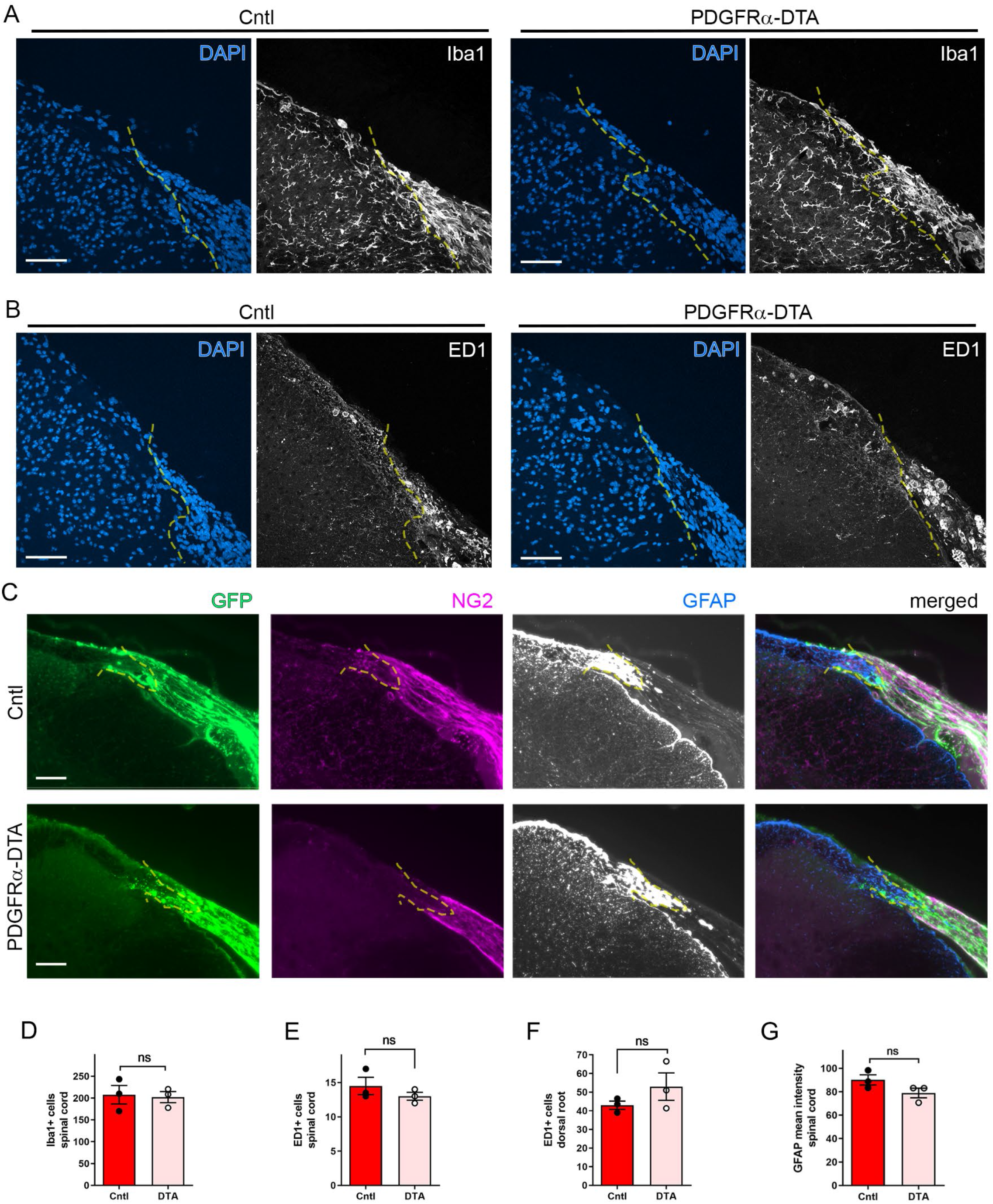
Microglia, macrophages and astrocytes in PDGFRα-DTA mice. **(A)** Transverse sections of a control and a PDGFRα-DTA mouse at 2 wpi, showing DAPI and Iba1-labeled microglia, and illustrating no noticeable differences in morphology or density of microglia in the spinal cord, DREZ, or DR of control and DTA mice. Scale bar: 100μm **(B)** Transverse sections from a control and PDGFRα-DTA mouse at 2 wpi, showing DAPI and ED1-labeled macrophages, and illustrating no noticeable differences in the distribution or density of macrophages in the spinal cord, DREZ, or DR of control and DTA mice. Scale bar: 100μm **(C)** Transverse sections of control and PDGFRα-DTA mice at 2 wpi, co-labeled for NG2 and GFAP, illustrating no noticeable differences in the density or distribution of astrocytes at the DREZ or within the spinal cord between control and DTA mice. Scale bar: 50μm **(D)** Quantitative comparisons of control and PDGFRα-DTA mice illustrating insignificant difference in the number of Iba1+ microglia in the spinal cord. Cntl, 207.8±21.20; DTA, 202.1±12.70. Unpaired Student’s t-test (n=3 mice/group). Error bars represent SEM. ns, p > 0.05 **(E, F)** Quantitative comparisons of control and PDGFRα-DTA mice, illustrating insignificant difference in the number of ED1+ macrophages in the spinal cord (E) or dorsal root (F). (E) Cntl, 14.50±1.258; DTA, 13.00±0.5774. (F) Cntl, 43.00±2.179; DTA, 52.94±7.330. Unpaired Student’s t-test (n=3 mice/group). Error bars represent SEM. ns, p > 0.05 **(G)** Quantitative comparisons of control and PDGFRα-DTA mice, illustrating insignificant difference in GFAP immunoreactivity in the spinal cord. Cntl, 90.10±4.363; DTA, 78.95±4.127. Unpaired Student’s t-test (n=3 mice/group). Error bars represent SEM. ns, p > 0.05

**Figure S6.**
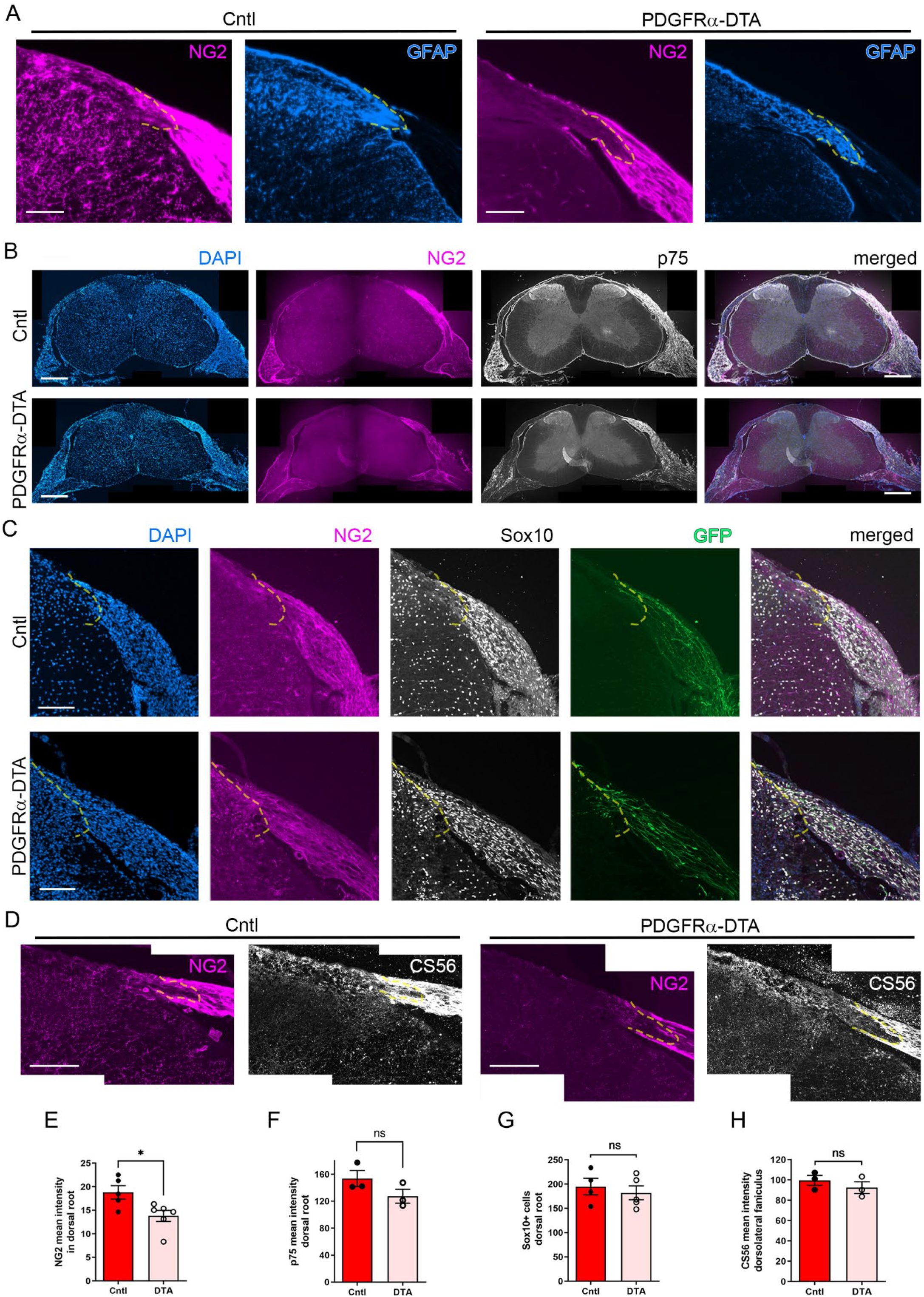
NG2/CSPG4, Schwann cells and CSPGs in PDGFRα-DTA mice. **(A)** Transverse sections of a control and a PDGFRα-DTA mouse with ∼80% OPCs ablated at 2 wpi, co-labeled for NG2 and GFAP, illustrating a marked reduction in OPCs and their processes in PDGFRα-DTA mouse. NG2 staining is slightly reduced peripherally along the DR of the PDGFRα-DTA mouse, as compared to control mouse. Scale bar: 100μm **(B)** Transverse sections of control and PDGFRα-DTA mice at 2 wpi, illustrating that p75 immunoreactivity in dorsal root of DTA mouse is comparable to that of control mouse. Scale bar: 500μm **(C)** Transverse sections of control and PDGFRα-DTA mice at 2 wpi, illustrating no noticeable difference in the number of Sox10-labeled Schwann cells present in the dorsal root. Scale bar: 100μm **(D)** Transverse sections of a control and a PDGFRα-DTA mouse, co-labeled for NG2 and for CS56 at 2 wpi, showing reduced NG2 but abundant CSPGs associated with the DREZ area of DTA mice. Scale bar: 100μm **(E)** Quantitative comparisons of control and PDGFRα-DTA mice, illustrating a slight reduction of NG2 immunoreactivity in dorsal roots of DTA mice. Cntl, 18.81±1.419; DTA, 13.81±1.173. Unpaired Student’s-t test (n=5 control, 6 DTA mice). Error bars represent SEM. *p ≤ 0.01 **(F)** Quantitative comparison of control and PDGFRα-DTA mice, illustrating insignificant difference in p75 immunoreactivity in the dorsal root. Cntl, 153.7±11.83; DTA, 117.5±18.55. Unpaired Student’s t-test (n=3 mice/group). Error bars represent SEM. ns, p > 0.05 **(G)** Quantitative comparison of control and PDGFRα-DTA mice, illustrating insignificant difference in the number of Schwann cells in the dorsal root. Cntl, 195.0±17.03; DTA 182.1±14.19. Unpaired Student’s t-test (n= 4 control, 5 DTA mice). Error bars represent SEM. ns, p > 0.05 **(H)** Quantitative comparisons of control and PDGFRα-DTA mice, illustrating insignificant difference in CS56 immunoreactivity measured in the dorsolateral funiculus including the DREZ. Cntl, 99.58.0±4.926; DTA 92.46±5.748. Unpaired Student’s t-test (n=3 control, 3 DTA mice). Error bars represent SEM. ns, p > 0.05

**Figure S7.**
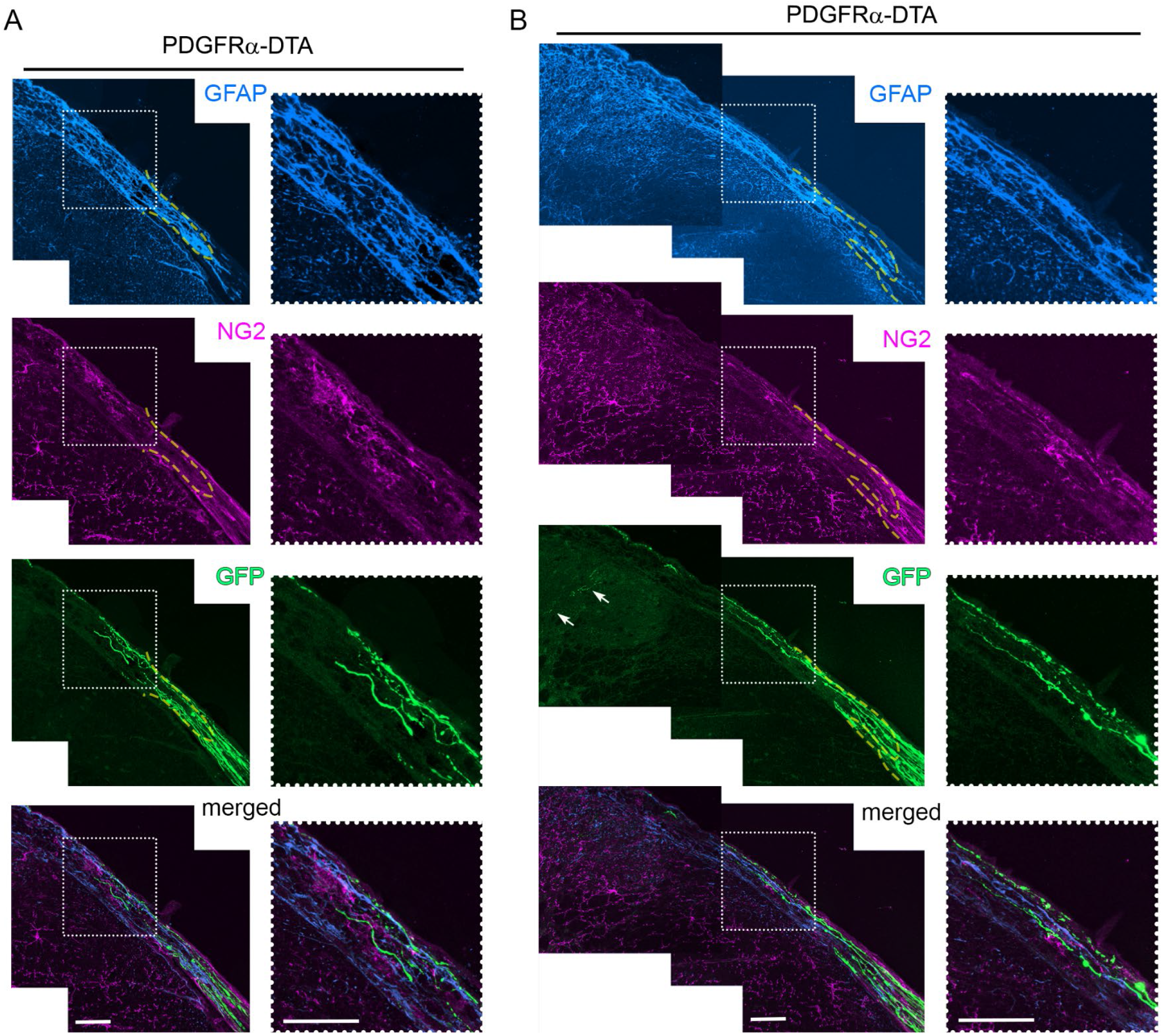
Axons regenerate deeper into the dorsal funiculus in PDGFRα-DTA mice at 4 wpi. **(A, B)** Two sets of transverse sections of PDGFRα-DTA mice at 4 wpi, co-labeled for GFAP and NG2, illustrating more numerous GFP+ axons (Green) that regenerated deeper into the dorsal funiculus than at 2 wpi (refer to Figure 7D). These axons terminate where OPCs are locally repopulated, in association with OPC processes (Magenta). Enlarged views of boxed area also indicate sparse distribution of OPCs where these axons regenerate. Arrows show axons reaching more deeply into the dorsal horn. Scale bar: 50μm

## STAR METHODS

### Key resources table

| REAGENT or RESOURCE | SOURCE | IDENTIFIER |
| --- | --- | --- |
| Antibodies |  |  |
| Rabbit polyclonal anti-GFAP | Dakocytomation | Cat# N1506; RRID: AB_10013482 |
| mouse monoclonal anti-GFAP | Sigma-Aldrich | Cat# G3893; RRID: AB_477010 |
| Goat polyclonal anti-MOG | R&D Systems | Cat# AF2439; RRID: AB_2145537 |
| Mouse monoclonal anti-SC2E | Cosmo Bio Co | Cat# CAC-GU01-M01AS-A; RRID: AB_1962608 |
| Goat polyclonal anti-p75NGFR | Neuromics | Cat# GT15057; RRID: AB_1611758 |
| Rat polyclonal anti-ED1 | AbD serotec | Cat# MCA1957; RRID: AB_322219 |
| Rabbit polyclonal anti-CGRP | Peninsula Lab | Cat# T-4032; RRID: AB_2307330 |
| Rabbit polyclonal anti-NF200 | Sigma-Aldrich | Cat# N4142; RRID: AB_477272 |
| Mouse monoclonal anti-SV2 | Developmental Studies Hybridoma Bank | Cat# SV2; RRID: AB_2315387 |
| Guinea pig polyclonal anti-NG2 | Dr. Dwight Bergles | N/A |
| Rat polyclonal anti-PDGFRα | BD Biosciences | Cat# 558774; RRID: AB_397117 |
| Rabbit-monoclonal anti-PDGFRα | abcam | *Cat# ab203491; RRID: AB_2892065* |
| Mouse monoclonal anti-PDGFRβ | abcam | *Cat# ab69506; RRID:AB_1269704* |
| Mouse monoclonal anti-APC-CC1 | Millipore | Cat# OP80; RRID: AB_213434 |
| Goat polyclonal anti-CD31/PECAM1 | R&D Systems | Cat# AF3628; RRID:AB_2161028 |
| Goat polyclonal anti-Sox10 | R&D Systems | Cat# AF2864; RRID:AB_442208 |
| Rabbit monoclonal anti-Vimentin | abcam | Cat# ab92547; RRID:AB_10562134 |
| Mouse monoclonal anti-CS56 | Sigma-Aldrich | Cat# C8035; RRID:AB_476879 |
| Rabbit polyclonal anti-GFP | abcam | Cat# ab290; RRID: AB_303395 |
| Rabbit polyclonal anti-olig2 | Millipore | Cat# AB9610; RRID: AB_570666 |
| Rabbit polyclonal anti-Ki67 | Millipore | Cat# AB9260; RRID: AB_2142366 |
| Rabbit polyclonal anti-NG2 IgG | Millipore | Cat# AB5320; RRID: AB_91789 |
| Mouse polyclonal anti-β-actin | Sigma-Aldrich | Cat# A5441; RRID: AB_476744 |
| Goat anti-mouse IgG (H+L), FITC | Invitrogen | Cat# A10530; RRID: AB_2534035 |
| Goat anti-mouse IgG (H+L), Alexa Fluor 568 | Invitrogen | Cat# A21124; RRID: AB_141611 |
| Goat anti-rabbit IgG (H+L), FITC | Invitrogen | Cat# A11034; RRID: AB_2576217 |
| Goat anti-rabbit IgG (H+L), Alexa Fluor 568 | Invitrogen | Cat# A11011; RRID: AB_143157 |
| Goat anti-rabbit IgG (H+L), Alexa Fluor 647 | Invitrogen | Cat# A21244; RRID: AB_141663 |
| Donkey anti-rabbit IgG (H+L), Alexa Fluor 546 | Invitrogen | Cat# A10040; RRID: AB_2534016 |
| Donkey anti-rabbit IgG (H+L), Alexa Fluor 647 | Invitrogen | Cat# A31573; RRID: AB_2536183 |
| Donkey anti-goat IgG (H+L), Alexa Fluor 488 | Invitrogen | Cat# A11055; RRID: AB_142672 |
| Donkey anti-rat IgG (H+L), TRITC | Jackson ImmunoResearch Laboratories | Cat# 712-025-150; RRID: AB_2340635 |
| Donkey anti-guinea pig IgG (H+L), Cy3 | Jackson ImmunoResearch Laboratories | Cat# 706-166-148; RRID: AB_23404061 |
| Donkey anti-guinea pig IgG (H+L), Dye 647 | Jackson ImmunoResearch Laboratories | Cat# 706-606-148; RRID: AB_2340477 |
| Goat anti-rabbit IgG (H+L), HRP | Santa Cruz | Cat# SC2030; RRID: AB_631747 |
| Goat anti-mouse IgG (H+L), HRP | Jackson ImmunoResearch Laboratories | Cat# 115-035-174; RRID: AB_2338512 |
| Goat anti-rat IgG (H+L), HRP | Jackson ImmunoResearch Laboratories | Cat# 112-035-003; RRID: AB_2338128 |
| Goat anti-guinea pig IgG (H+L), 10nm gold particles | EMS | Cat# 25329 |
| Bacterial and virus strains |  |  |
| scAAV2-eGFP | Dr. George M Smith | N/A |
| Chemicals, peptides, and recombinant proteins |  |  |
| Ketamine | Covetrus | Cat# 11695-0703-1 |
| Xylazine | Akom, Inc. | Cat# 59399-110-20 |
| Buprenorphine | Par pharmaceuticals | Cat# 42023-0179-05 |
| Euthasol | Virbac | Cat# 051311-050-01 |
| paraformaldehyde | EMS | Cat# 157-8 |
| glutaraldehyde | EMS | Cat# 16320 |
| 0.1M phosphate buffer | EMS | Cat# 19340-72 |
| 4-hydroxytamoxifen | Sigma-Aldrich | Cat# H6278 |
| DiI | invitrogen | Cat# N22880 |
| Kwik-Sil | World Precision Instruments, Inc. | Kwik-sil |
| Epredia™ M-1 Embedding Matrix | Thermo Scientific | Cat# 1310 |
| Vectorshield | Vector Laboratories | Cat# H-1000 |
| Glycine powder | Sigma-Aldrich | Cat# 50046 |
| Bovine Serum Albumin | Sigma-Aldrich | Cat# A3059 |
| Triton X-100 | Sigma-Aldrich | Cat# T8787 |
| Osmium tetroxide | EMS | Cat# 19150 |
| Araldite 6005 | EMS | Cat# 10920 |
| Uranly acetate | EMS | Cat# 22400 |
| Sodium Citrate | EMS | Cat# 21140 |
| Toluidine Blue O | EMS | Cat# 22050 |
| Borax | EMS | Cat# 21130 |
| photo-initiator 2,20-Azobis[2-(2-imidazolin-2-yl) propane] dihydrochloride | Wako Chemicals USA | Cat# VA-044 |
| acrylamide | Bio-Rad | Cat# 161-0140 |
| Bis-acrylamide | Bio-Rad | Cat# 161-0142 |
| Boric acid | Sigma-Aldrich | Cat# B7901 |
| SDS | Sigma-Aldrich | Cat# L337 |
| FocusClear | CelExplorer | Cat# FC-1010 |
| ECL Plus | GE Healthcare | Cat# RPN2236 |
| Anti-BSC1 Griffonia Simplificonia (BSL1) | Vector Laboratories | Cat# L1100; RRID: AB_2336491 |
| DMEM | Gibco | Cat# 11960-044 |
| B27 supplement | Gibco | Cat# 17504-044 |
| sodium pyruvate | Gibco | Cat# 11360-070 |
| insulin | Sigma-Aldrich | Cat# I6634 |
| Apo-transferrin | Sigma-Aldrich | Cat# T1147 |
| bovine serum albumin | Sigma-Aldrich | Cat# A4161 |
| progesterone | Sigma-Aldrich | Cat# P8783 |
| putrescine | Sigma-Aldrich | Cat# 5780 |
| sodium selenite | Sigma-Aldrich | Cat# S5261 |
| N-acetyl-L-cysteine (NAC) | Sigma-Aldrich | Cat# C7477 |
| PDGF | Sigma-Aldrich | Cat# SRP3228 |
| bFGF | Sigma-Aldrich | Cat# SRP4038 |
| Hank's balanced salt solution (HBSS) | Sigma-Aldrich | Cat# H6648 |
| dispase II | Roche Applied Sciences | Cat# 04942078001 |
| HEPES-buffered saline | Sigma-Aldrich | Cat# 51558 |
| Percoll gradient | GE Healthcare | Cat# 17-0891-01 |
| fura-2AM | Thermo Scientific | Cat# J62728.MCR |
| **Experimental models: Organisms/strains** |  |  |
| Mouse: Thy1-YHFM |  |  |
| Mouse: Thy1-YHFH |  |  |
| Mouse: PDGFRα-creER<sup>TM</sup> | Jackson Laboratory | RRID: IMSR_JAX:018280 |
| Mouse: Rosa-DTA | Jackson Laboratory | RRID: IMSR_JAX:009669 |
| Mouse: Pngfα-CreER<sup>T2</sup> | Jackson Laboratory | RRID:IMSR_JAX:032770 |
| Mouse: Ai14 | Jackson Laboratory | RRID:IMSR_JAX:007914 |
| **Software and algorithms** |  |  |
| MetaFluor Fluorescence Ratio Imaging software | Molecular devices | RRID: SCR_014294 |
| AxioVision Imaging System | Zeiss | RRID: SCR_002677 |
| Leica LAS AF software | Leica | RRID: SCR_013673 |
| Imaris | Bitplane | RRID: SCR_007370 |
| Photoshop | Adobe | RRID: SCR_014199 |
| Prism 9 | GraphPad | RRID: SCR_002798 |
| ImageJ | NIH | RRID: SCR_003070 |
| Fiji | NIH | RRID: SCR_002285 |
| **Other** |  |  |
| Superfrost Plus microscope slides | Fisher Scientific | Cat# 22-230-892 |
| Leica TCS SP5 II multi-photon confocal microscope | Leica Microsystems | N/A |
| Leica SP8 confocal microscope | Leica Microsystems | N/A |
| epifluorescence M2 microscopy | Zeiss | N/A |
| Leica Ultracet R microtome | Leica Microsystems | N/A |
| Leica cryostat | Leica Microsystems | Cat# CM3050S |
| Diamond knife (Ultra 45°, 4mm) | Diatom | N/A |
| Biobrane | UDL Laboratories, Inc. | Cat# 8459-0095-25 |
| Gore Preclude MVP Dura Substitute | Gore and Associates | Cat# 1MVP401 |
| 6-0 suture | Myco medical | Cat# SK681-SI |

### Resource availability Lead contact

Further information and requests for resources and reagents should be directed to and will be fulfilled by the lead contact, Young-Jin Son.

### Materials availability

All reagents generated in this study are available from the lead contact without restriction.

### Data and code availability

- All data reported in this paper will be shared by the lead contact upon request.
- This paper does not report the original code.
- Any additional information required to realize the data reported in this paper is available from the lead contact upon request.

### Experimental model and subject details

#### Animals

All animal care and procedures were conducted in accordance with the National Institute of Health guidelines and with approval from the Institutional Animal Care and Use Committee at Lewis Katz School of Medicine at Temple University. For *in vivo* imaging or CLARITY tissue clearing, C57BL/6J or the transgenic strains, *Tg(Thy1-EGFP)MJrs/J* (Thy1-GFP-M; JAX 007788) or *B6.Cg-Tg(Thy1-YFP)HJrs* (Thy1-YFP-H; JAX 003782) were purchased from Jackson Laboratory. For OPC ablation, *B6N.Cg-Tg(Pdgfra-cre/ERT)467Dbe/J* (PDGFRα-creER^TM^; JAX 018280) and *B6.129P2-Gt(ROSA) 26Sor^tm1(DTA)Lky/J^* (Rosa26-DTa; JAX 009669) purchased from Jackson Laboratory were crossed to generate *PDGFRα-creER^TM^; Rosa26-DTa* mice. *B6.129S-Pdgfrα^tm1.1(cre/ERT2)Blh^*/J (PDGFRα-CreER^T2^; JAX:032770) and *B6.Cg-Gt(ROSA)26Sor^tm14(CAG-tdTomato)Hze^/J* (Ai14; JAX:007914) from Jackson Laboratory were crossed to generate PDGFRα-tdTomato reporter mice. Adult mice of either sex, 2-3 months old, were used for all experiments. Age matched *PDGFRα-creER^TM^* or *Rosa26-DTa* mice were used as the control.

### Method details

#### Dorsal root crush

For unilateral cervical (C5-C8) or lumbar (L4 and L5) root crush, mice were anesthetized with an intraperitoneal injection of xylazine (8 mg/kg) and ketamine (120 mg/kg). Following a 2- to 3- cm-long incision in the skin of the back, the spinal musculature was reflected and the C5-C8 or L3–S1 spinal cord segments exposed by hemilaminectomies. A small incision was made in the dura overlying each targeted DR. A fine forceps (Dumont #5; Fine Science Tools, Foster City, CA) was introduced subdurally and each DR was crushed for 10 s. To avoid scar formation and possible compression, we applied a piece of thin silastic membrane (Biobrane, Bertek Pharmaceuticals, Sugarland, TX) over the hemilaminectomy site and covered it with a layer of thicker artificial dura (Gore Preclude MVP Dura Substitute, W.L. Gore and Associates, Flagstaff, AZ). The overlying musculature was closed with 6-0 sutures, and the skin closed with wound clips. All animals received subcutaneous injections of saline (0.9% w/v NaCl) and buprenorphine (0.05 mg/kg) for postoperative pain management.

### Intraganglionic injection of AAV2-GFP

Recombinant adeno-associated virus 2 carrying eGFP (AAV2-GFP) with CMV-enhanced chicken β-actin (CAG) promoter was purchased from Vector Biolabs (Malvern, PA; stock #7072) or generated as previously described (Liu et al., 2014). AAV2-GFP was microinjected into a DRG using a micropipette pulled to a diameter of 0.05 mm and a nanoinjector (B203XVY Nanoliter 2000; World Precision Instruments, Sarasota, FL). For each injection, the micropipette was introduced 0.5 mm into the DRG and a total volume of AAV (1 μl/DRG) containing >1 X 10^13^ GC/ml injected over a 10 min period. The glass needle was left in place for 5 min after each injection.

### 4-HT administration

For OPC ablation, 4-hydroxytamoxifen (4-HT; Sigma # H7904, St. Louis, MO) was injected into 2-3 months old *PDGFRα-CreERTM;ROSA26-iDTa* mice. A 20 mg/ml stock of 4-HT in 100% ethanol was prepared by sonication, and a 1 mg aliquot mixed 1:5 with sunflower oil. After evaporation of the ethanol by speedvac, 1 mg of 4-HT was injected intraperitoneally once daily for 5 consecutive days beginning 2 dpi.

### CLARITY tissue clearing

Longitudinal slices of the dorsal spinal cord (<1mm thick) were optically cleared using the CLARITY protocol modified for passive clearing (Tomer et al., 2014). Adult mice that received unilateral L4 and L5 root crush, followed by AAV2-GFP intraganglionic injections, were perfused at 2 wpi with 20 ml of ice-cold 0.1 M PBS, followed by 20 ml of ice-cold hydrogel monomer solution [4% acrylamide (Bio-Rad, #161-0140), 0.25% azo-initiator (Wako, #VA-044), 4% Paraformaldehyde (Electron Microscopy Sciences, #15714-S) in PBS (Invitrogen, #70011-044)]. Each spinal cord was quickly dissected, its dorsal surface longitudinally sliced with attached dorsal roots, and the dorsal spinal cord placed in 20 ml of cold hydrogel in a 50 ml conical tube. After 24 hours at 4oC, the tube was placed in a desiccation chamber with its lid twisted open but not removed. The desiccation chamber was attached to a vacuum for 10 minutes, the chamber then purged with nitrogen gas, and the tube lid quickly screwed shut to prevent exposure to the outside air. The 50 ml tubes were placed in a 37°C water bath for 2 hours to allow for polymerization of the hydrogel and tissue. After polymerization, spinal cord was transferred into a 50 ml tube with clearing solution 20 mM lithium hydroxide monohydrate (Sigma-Aldrich, #254274), 200 mM boric acid (Sigma-Aldrich, #B7901), 4% SDS (Sigma-Aldrich, #L3771, pH to 8.5] and placed in an incubator with agitation at 37°C. Every 24 hours the clearing solution was replaced with fresh clearing solution for ∼ 7 days or until the tissue had completely cleared. For microscopy, tissue was submerged in FocusClear (CelExplorer Labs Co., #FC-101) for 24 hours at 37°C to allow for refractive index matching. Tissue was then placed in an imaging well constructed on a standard microscope slide with a thin barrier of BluTack putty (Bostick, Fort Wayne, IN) sealed with a Willco-Dish (Ted Pella, #14032-120) and Kwik-Sil (World Precision Instruments, #KWIK-SIL).

### Two-photon and widefield *in vivo* imaging

Both widefield and two-photon microscope set-ups were used for repeated imaging of DR axons at the DREZ in living mice, as we previously described (Skuba *et al.*, 2014). Widefield imaging was performed using a fluorescence stereomicroscope (M205C, Leica Microsystems) with a fast shutter and a CCD camera (ORCA-Rx2, Hamamatsu) controlled by MetaMorph software (Molecular Devices). Images were acquired either as single snapshots or as multiple streams of 10–20 frames acquired within 30–40 msec. exposure time. In-801 focus images were then selected, and an overview montage was created using Photoshop (Adobe Systems). A temperature controller (ATC 1000, World Precision Instruments, Inc.) and attached heating pad maintained the mouse’s body temperature. For prolonged (>1 hr.) sessions, continuous inhalation anesthesia was consistently delivered via a laboratory animal anesthesia system (V-1 LAAS, VetEquip). 2-photon imaging was performed using an upright microscope optimized for confocal and multiphoton microscopy (TCS SP5 II, Leica Microsystems). After the mouse was mounted to a stereotaxic holder (SR-AM and STS-A, Narishige), imaging was performed with a Spectra Physics MaiTai laser tuned to ∼890 nm for 2-photon excitation of YFP. Image acquisition used a 20X 0.7 NA water immersion lens controlled by Leica LAS AF software. For an axon ending of interest, a Z-stack was taken with line average every 1 μm throughout the entire focal area spanning 20-50 μm. Laser power and offset were adjusted to reduce background noise. If necessary, alignment software (Auto Aligner, Bitplane) was used to manually align off-centered optical sections. Maximum intensity projections along the Z-axis are presented as the final images.

### Immunohistochemistry

Two or four weeks after DR crush, mice were sacrificed by an overdose of Euthasol and perfused transcardially with 0.9% saline followed by 4% paraformaldehyde (PFA) in 0.1 M PBS. The spinal cords with attached DRs and DRGs were removed and examined in wholemounts to exclude those tissues with poor transduction of AAV2-GFP and/or spared axons. Tissues were then processed for analysis on cryostat sections or as wholemounts. For analysis on cryostat sections, spinal cords were post-fixed in 4% PFA overnight at 4°C and cryoprotected in 30% sucrose for 24 hr. The tissues were then embedded in M-1 Embedding Matrix (Thermo Fisher Scientific, Waltham, MA), transversely sectioned at 20μm using a cryostat (Leica Microsystems, Germany), and mounted directly on slides (Superfrost Plus, Fisher Scientific, Pittsburgh, PA).

For immunolabeling, sections were rinsed in PBS for 30 min. followed by 10 min. incubation with 0.1 M glycine in PBS, and 15 min. incubation with 0.2% Triton X-100, 2% bovine serum albumin (BSA) in PBS (TBP). Sections were incubated with primary antibodies overnight at 4°C, washed three times for 30 min. with 2% BSA in PBS, and incubated with secondary antibodies for 1 hr. at room temperature. After rinsing in PBS, sections were mounted in Vectashield (Vector Laboratories, Burlingame, CA) and stored 832 at –20°C until examination. For wholemount immunostaining of spinal cords, spinal cords with attached DRs were post-fixed in 4% PFA for 2 hr. at 4°C and the dura mater removed. The spinal cord was then rinsed with PBS for 30 min., incubated for 10 min. in 0.1 M glycine in 2% BSA/PBS, and permeabilized with pre-cooled methanol at –20°C for 10 min. After extensive rinsing in PBS, the spinal cord was incubated with primary antibody diluted in TBP at room temperature overnight, rinsed thoroughly in TBP the following day, and incubated with secondary antibodies for 2 hr. at room temperature. After rinsing in PBS, a thin slice of dorsal spinal cord (∼2 mm thick) with attached DR stumps was cut with a microscissors and mounted on slides in Vectashield (Vector Laboratories).

### Transmission and immunoelectron microscopy

For transmission electron microscopy, mice were perfused transcardially with 2% paraformaldehyde and 2.5% glutaraldehyde in 0.1 M Na-cacodylate buffer. Lumbar spinal cord segments (L3-L6) with attached DRs were removed as one piece, mounted on agarose support, and placed in the vibratome well. The most superficial longitudinal slice containing the DREZ (<250 μm thickness) was cut and further processed for TEM, as previously described (Di Maio *et al.*, 2011). For immunogold labeling of OPCs, mice were perfused with ice-cold 0.9% heparinized saline followed by 4% paraformaldehyde and 0.2% glutaraldehyde in 0.1M PBS (pH 7.4). After 2 hr. postfixation in the same fixative at 4°C, the spinal cord segment was embedded in 10% gelatin, cooled to 4°C, and a vibratome section of superficial spinal cord segment (∼80 mm thickness) collected in PBS. The vibratome sections were then incubated with 1% sodium borohydride in PBS for 30 min., followed by permeabilization with 0.1% Triton-X-100 in PBS for 20 min. at 4°C. The sections were blocked with blocking solution (1% BSA, 3% NGS, 0.04% Triton-X-100 in PBS) for 30 min. and incubated with guinea pig anti-NG2 antibody (1:500, a gift from Dr. Dwight Bergles, Johns Hopkins University) diluted with blocking buffer for 24-48 hr. at 4°C. After rinsing, sections were incubated with 10 nm gold-conjugated goat anti-guinea pig IgG (1:200, Electron Microscopy Sciences, #25329) in blocking buffer for 3 hr. at room temperature. After rinsing, sections were post-fixed in 2% glutaraldehyde in PBS for 10 min.

Silver enhancement was then performed in the dark with the HQ Silver Kit (Nanoprobes, NY) as directed by the manufacturer. Sections were post-fixed again with 1% osmium tetroxide in PBS for 10 min. and then incubated with 2% uranyl acetate in ddH2O 863 for 10 min. Sections were then dehydrated in an ascending ethanol series, followed by acetone series, and finally infiltrated with Araldite 502 Embed 812 resin. After polymerization, serial ultrathin sections (60–70 nm) were cut and mounted on Pioloform-coated slot grids. They were observed with a JEOL 1010 electron microscope at the Electron Microscopy Resource Lab, University of Pennsylvania.

### Western blotting

The dura was removed from the spinal cord, which was then snap-frozen in liquid nitrogen and stored at -80oC. Spinal cords were lysed in RIPA buffer (25 mM Tris pH 7.5, 150 mM NaCl, 1% Triton X-100, 0.5% sodium deoxycholate, 1 mM EDTA, 0.1% SDS). Lysis was for 40 min. on ice, followed by microcentrifugation at 14,000 rpm for 20 min. Protein concentration was determined using the BCA assay (ThermoFisher). Primary antibodies were used at the following concentrations for Western blots: rabbit anti-NG2 (Millipore, AB5320; 1:500), mouse anti-β-actin (Sigma, A5441; 1:1,000), and rat anti-PDGFRα (BD Biosciences, 558774; 1:500).

Secondary antibodies for Western blots were HRP-conjugated goat anti-rabbit antibody (Santa Cruz, SC2030; 1:10,000), HRP-conjugated goat anti-mouse antibody (Jackson ImmunoResearch Laboratories, 115-035-174, 1:10,000), or HRP-conjugated goat anti-rat antibody (Jackson ImmunoResearch Laboratories, 112-035-003; 1:10,000). Blots were developed using ECL, ECL Plus (GE Healthcare, RPN2236).

### Antibodies for immunohistochemistry

Primary antibodies were used at the following concentrations for immunohistochemistry: rabbit anti-glial fibrillary acidic protein (GFAP, N1506; 1:500), mouse anti-GFAP (Sigma-Aldrich, G3893; 1:500), goat anti-myelin oligodendrocyte glycoprotein (MOG; R&D Systems, AF2439; 1:200), mouse anti-SC2E (Cosmo Bio Co., CAC-GU01-M01AS-A; 1:10,000), goat anti-p75NGFR (Neuromics, GT15057; 1:500), rat anti-ED1 (AbD serotec, MCA1957; 1:400), rabbit anti-CGRP (Peninsula Lab., T-4032; 1:2,000), rabbit anti-NF200 (Sigma-Aldrich, N4142, 1:500), mouse anti-SV2 (Developmental Studies Hybridoma Bank, SV2; 1:10), guinea pig anti-NG2 (1:500, a gift from Dr. Dwight Bergles, Johns Hopkins University), rat anti-PDGFRα (BD Biosciences, 558774; 1:500), mouse anti adenomatous polyposis coli antigen (APC; clone CC1; Millipore, OP80-100UG; 1:100), rabbit anti-PDGFRα (abcam, ab203491; 1:500), mouse anti-PDGFRβ (abcam, ab69506; 1:500), mouse anti-CS56 (Sigma-Aldrich, C8035 1:1,000), goat anti-CD31/PECAM (R&D Systems, AF3628; 1:200), rabbit anti-vimentin (abcam, ab92547;

1:1,000). Secondary antibodies were FITC conjugated goat anti-mouse IgG (Invitrogen, A10530; 1:400), Alexa-Fluor 568-conjugated goat anti-894 mouse IgG (Invitrogen, A21124; 1:400), Alexa-Fluor 488-conjugated goat anti-rabbit IgG (Invitrogen, A11034; 1:400), Alexa-Flour 568 conjugated goat anti-rabbit IgG (Invitrogen, A11011; 1:400), Alexa-Flour 546 conjugated donkey anti-rabbit IgG (Invitrogen, A10040; 1:400), Alexa-Fluor 647-conjugated donkey anti rabbit IgG (Invitrogen, A31573; 1:400), Alexa-Fluor 647-conjugated goat anti-rabbit IgG (Invitrogen, A21244; 1:400), Alexa-Flour 488 conjugated donkey anti-goat IgG (Invitrogen, A11055; 1:400), rhodamine (TRITC)-conjugated donkey anti-rat IgG (Jackson ImmunoResearch Laboratories, 712-025-150; 1:400), Cy3-conjugated donkey anti-guinea pig IgG (Jackson ImmunoResearch Laboratories, 706-166-148; 1:400), and Dye 647-conjugated donkey anti guinea pig IgG (Jackson ImmunoResearch Laboratories, 706-606-148; 1:400).

### OPC purification and co-culture with Percoll-purified DRG neurons

OPCs were isolated from P16-P18 C57BL/6 mice by immunopanning as previously described (Cahoy et al., 2008). Dissociated spinal cords were resuspended in panning buffer. To deplete microglia, the cell suspension was sequentially panned on two BSL1 panning plates (Vector Laboratories, #L1100). To purify OPCs, the cell suspension was then incubated on two PDGFRα plates coated with rat anti-PDGFRα IgG (BD Biosciences, 558774; 1:500). The adherent cells were washed with PBS to remove all antigen-negative nonadherent cells. The adherent cells were trypsinized and plated onto poly-D-lysine (PDL)-coated plates. The cultures were maintained under proliferating conditions by the addition of PDGFAA (Sigma, SRP3228; 10ng/ml) and bFGF (Sigma, SRP4038; 20 ng/ml) to the serum-free medium (DMEM, B27 supplement, 1X Sato supplement, 1X Trace Elements B, 10 ng/ml d-biotin, 2 mM Glutamine, 1 mM sodium pyruvate, 5 ug/ml insulin, 5 μg/ml N-acetyl-L-cysteine, 500 mM Forskolin, 100 U/ml Penicillin streptomycin). Most cells (>95%) expressed OPC markers Olig2, NG2, and PDGFRα. After 5 days of incubation, percoll-purified DRG neurons were applied to cultured OPCs and cocultured for 5 days. For isolating DRG neurons, ∼20 DRGs were removed from adult C57BL/6 mice and then dissociated in 1 ug/ml collagenase A and 2.4 U/ml dispase II. The dissociated DRGs were added to a two-layered Percoll gradient (12.5% Percoll layered over 28% Percoll solution with DMEM) and then subjected to density-gradient centrifugation. The interphase fraction was subsequently removed, spun, and resuspended in 20 μl of DMEM. A small amount of the cell suspension (<5 μl) was added sparingly to cultured OPCs in the OPC proliferating medium. The cocultures were incubated in 5% CO2 at 37oC for 5 days.

### Calcium imaging

Calcium imaging was performed using fura-2-based microfluorimetry and imaging analysis as we previously described (Dou et al., 2018). OPC-DRG neuron cocultures on coverslips were loaded with 4 μM of fura-2AM (Life Technologies, Grand Island, NY) for 30 min. at room temperature in HBSS, washed, and further incubated in normal bath solution (Tyrode’s; 140 mM NaCl, 5 KCl mM, 2 CaCl2 mM, 1 MgCl2 mM, 10 HEPES mM, and 5.6 mM glucose).

Coverslips were mounted in a small perfusion chamber (Warner Instruments, Model RC-25) and continuously perfused at 5–7 ml/min with Tyrode’s solution. 100 nM capsaicin in Tyrode’s solution, with or without 20μM CNQX and 10μM AP5, was locally infused for 20 sec., using a glass electrode (∼500 μm ID) positioned ∼1.0 mm away from cells. Images were acquired at 3 s. intervals at room temperature using an Olympus inverted microscope equipped with a CCD camera (Hamamatsu, ORCA-03G). The fluorescence images were recorded and analyzed using the software MetaFluor 7.7.9 (Molecular Devices). The fluorescence ratio was determined as the fluorescence intensities excited at 340 and 380 nm with background subtraction.

### Quantification and statistical analysis

#### Fluorescence microscopy, cell counting, and densitometric analysis

Z-stack fluorescent images were acquired on a Zeiss Axio Imager widefield microscope or on a Leica SP8 confocal microscope from 20μm serial transverse sections of spinal cords. Acquisition parameters were kept constant between different groups. Acquired images were merged using Imaris (Bitplane, Windsor, CT) and processed in Adobe Photoshop with minimal manipulations of brightness and contrast (Adobe Inc, San Jose, CA). For counting Ki67, Olig2, Iba1, ED1, and Sox10+ cells, 3 to 6 representative, non-adjacent sections taken from spinal cord segments of each mouse (n=3-6 per cohort) were carefully selected. Cell counting was performed blindly using ImageJ/Fiji (NIH). Immunoreactive cells were manually counted in ipsilateral regions of dorsal lateral funiculus, dorsal root, or dorsal spinal cord, and then averaged by the number of evaluated sections and animals. For comparative analysis of OPC ablation, CSPG deletion, GFAP, and p75 immunoreactivity, images were converted to grayscale, subthresholded to the background, and calculated using ImageJ/Fiji (NIH). Intensity was defined as the average gray value obtained from all pixels within the region of interest 955 above background. Data are presented as the mean fluorescence intensities (MFI) in each region of interest, calculated by subtracting the mean value of randomly selected background regions lacking immunoreactivity of interest.

### Analysis of axon regeneration across the DREZ

The DREZ was defined as the ∼100μm wide CNS territory beyond the boundary between the CNS and PNS. The yellow dotted lines (e.g., Figure 3A) denote the boundary (termed also borderline or border), demarcated by GFAP immunolabeling of astrocytic processes that extend peripherally. When GFAP was not immunolabeled, the borderline was identified based on the greater abundance of cell nuclei in the PNS than CNS (Zhai *et al.*, 2021). For comparative evaluation of regeneration, we considered that an axon penetrated the DREZ when it extended at least 100μm beyond the borderline. For quantitative analysis, digital images were captured from 3 to 6 representative, non-adjacent sections taken from spinal cord segments of each mouse (n = 3–6 mice per cohort). A raw image was converted to a binary image using ImageJ with a threshold that appropriately separated GFP and background fluorescence. Lines were drawn at 100μm before the borderline in the PNS, at the borderline, and at 100μm intervals into CNS territory. The number of intersections of GFP+ axons at these distances was counted. The number of axons that crossed the border was normalized by the number of GFP+ axons counted at 100μm before the borderline in the PNS, and then averaged by the number of evaluated sections and animals. This quantification resulted in the ‘axon number index’ that indicated the relative number and distance of axons that regenerated into the CNS or spinal cord.

### Statistical analysis

All statistical analyses were performed using PRISM 9.0 (GraphPad). Statistical analysis was done using the two-sample Mann-Whitney test or unpaired t-test for two-group comparisons, and analysis of variance (ANOVA) with Tukey’s or Sidak’s test for multiple comparisons, as appropriate. All data were presented as mean ± SEM or SD as indicated in the figure legend. A p-value of 0.05 or less was considered statistically significant.

## SUPPLEMENTAL INFORMATION

Supplemental figures and figure legends – 7 Supplemental figures Supplemental experimental procedures - TBD

